# Deciphering of single-cell chromatin accessibility and transcriptome reveals the discrepancy for *ex vivo* human erythropoiesis

**DOI:** 10.1101/2025.04.15.648947

**Authors:** Zijuan Xin, Wei Zhang, Lijuan Zhang, Guangmin Zheng, Tingting Jin, Meng Li, Yangliu Shao, Liping Dou, Shengwen Huang, Zhaojun Zhang, Xiangdong Fang

**Author notes:** Correspondence: Xiangdong Fang, Zhaojun Zhang. These authors contributed equally (Zijuan Xin, Wei Zhang, Lijuan Zhang, Guangmin Zheng). E-mail Address: Zijuan Xin, Wei Zhang, Lijuan zhang, Guangmin Zheng, Tingting Jin, Meng Li, Yangliu Shao, Liping Dou, Shengwen Huang, Zhaojun Zhang, Xiangdong Fang.

## Abstract

**Background:** Erythroid cells can be generated from hematopoietic stem and progenitor cells (HSPC) derived from various sources; however, few studies decode *ex vivo* human erythropoiesis for the discrepancies at single-cell multi-omics resolution and uncover the underlying restraints for erythrocyte regeneration.

**Results:** We deciphered *ex vivo* human erythropoiesis at two differentiation states from three sources at single-cell chromatin accessibility and transcriptomes level. We identified detrimental myeloid differentiation tendencies during early differentiation. These tendencies were linked to low glutamine activity in cord blood- and iPSC-derived erythropoiesis. The erythroid progenitor differentiation is restricted by cell cycle and hypoxia signaling deficiencies, which are pronounced in the iPSC-derived erythropoiesis. We delineated the erythroid differentiation trajectory of various *ex vivo* erythropoiesis systems, and revealed a distinct roadmap from HSC to orthochromatic erythroblast with unprecedented resolution. The integrative analysis of single-cell chromatin accessibility and transcriptome across developmental stages uncovered a dynamic coordination, and highlighted the pivotal role of chromatin accessibility and associated enhancers in regulating *ex vivo* erythropoiesis. Cell–cell communications in the *ex vivo* erythropoiesis system were not as well established as those in the BM, suggesting that modulating cell-cell communication signals in distinct *ex vivo* erythropoiesis system may facilitate erythrocyte regeneration.

**Conclusions:** This study comprehensively characterized the discrepancies and constraints in *ex vivo* human erythropoiesis at single-cell multi-omics resolution, offering novel strategies to overcome these constraints. These insights are critical for advancing functional erythrocyte generation and have significant implications for clinical applications.

## Background

Despite the annual collection of over 100 million units of blood donations worldwide, a persistent scarcity emerges in the face of pivotal global trends. *In vitro* production of red blood cells (RBCs) offers a solution to the worldwide shortage of blood for transfusion purposes. Erythroid cells can be regenerated *in vitro* from hematopoietic stem progenitor cells (HSPC) derived from various origins, including pluripotent stem cells (PSC), cord blood (CB), peripheral blood (PB), and bone marrow (BM).[1] Human PSC are ideal and theoretically infinite resources for erythrocyte generation. However, PSC-derived erythrocytes have unsatisfactorily low adult-type globin expression and extremely low enucleation rate.[2, 3] CB- and PB-derived erythrocytes exhibit higher adult globin expression and enucleation compared to those from PSC. However, adult PB progenitors exhibit a more constrained ability to proliferate, making them less ideal for the mass production of RBCs needed for therapeutic applications.[4] CB-derived progenitors demonstrate a higher potential for expansion than those from PB; however, the erythroid cells produced are limited, and these cells tend to exhibit characteristics of a fetal phenotype rather than those of mature cells.[5] The erythroid cells derived from these sources demonstrate differential erythroid maturity and regeneration efficiency. Currently, few studies decode *ex vivo* human erythropoiesis for the discrepancies at single-cell multi-omics resolution and uncover the underlying restraints for erythrocyte regeneration.

Erythropoiesis encompasses early erythropoiesis and terminal erythroid differentiation. These processes are regulated at multiple levels, including gene expression regulation, by coordinating chromatin accessibility dynamics.[6] A unique feature of erythrocyte regeneration is that each cell division is accompanied by cell differentiation, during which daughter cells differ in structure and function from mother cells,[7] yet traditional large-scale omics techniques cannot accurately reveal these dynamic, cell-type specific changes in gene expression and chromatin accessibility, especially for those uncertain cell types in the differentiation system, or the erythroid differentiation states and dynamics in high-resolution during *ex vivo* erythropoiesis.[2, 6] Recently, single-cell studies have elucidated regulators in hematopoiesis[8–10], and *ex vivo* erythropoiesis systems established in our laboratory.[5, 11] However, the absence of comprehensive single-cell omics data from *ex vivo* erythropoiesis systems limits our ability to compare erythroid differentiation discrepancies and uncover the constraints affecting *ex vivo* erythropoiesis. Addressing these gaps is crucial for optimizing the efficient generation of functional RBCs.

In this study, we employed single-cell transcriptome and chromatin accessibility sequencing for continuously differentiated erythroid cells derived from human cord blood, peripheral blood, and induced pluripotent stem cells (iPSC). We comprehensively elucidated the discrepancies and underlying constraints in the *ex vivo* human erythropoiesis systems, offering a practical strategies and resources for optimizing erythrocyte regeneration.

## Results

### Single-cell omics profiling of *ex vivo* human erythropoiesis

To explore the discrepancies and constraints in *ex vivo* human erythropoiesis, we collected continuously differentiated erythroid cells on days 7 and 14 derived from three human sources, namely cord blood (CB), peripheral blood (PB), and iPSC (SC), and conducted single-cell chromatin accessibility and transcriptome sequencing (Figure 1A). After rigorous data quality control, 99,776 cells from three *ex vivo* erythropoiesis systems in two differentiation states were pooled for downstream analysis (Additional file 1: Figure S1A–C). The semi-supervised cell automation annotation tool SCINA combined with cell clustering analysis showed 10 cell types with similar distribution patterns between transcriptome and chromatin accessibility levels (Figure 1B, Additional file 2: Table S1). The cell annotation results revealed that the highest proportion (55.6%) was erythroid cells, including erythroid progenitors (ErP, 29.4%) and erythroblasts (26.2%). In addition, a large number of myeloid lineages including macrophages (14.1%) and granulocyte-macrophage progenitors (GMP, 10.4%), hematopoietic stem cells and multipotent progenitors (HSC/MPP, 11.6%), and other hematopoietic cells were also observed in the three *ex vivo* erythropoiesis systems (Figure 1C), consistent with previous findings. [5, 12, 13] A notable enrichment of erythroid cells and cell type dynamic changes were observed on day 14 compared with that on day 7 (Figure 1D and 1E, Additional file 1: Figure S1D). Furthermore, genes associated with globin synthesis and oxygen transport showed significantly elevated expression and enhanced accessibility during erythroid differentiation (Additional file 1: Figure S1E-F).

**Figure 1.**
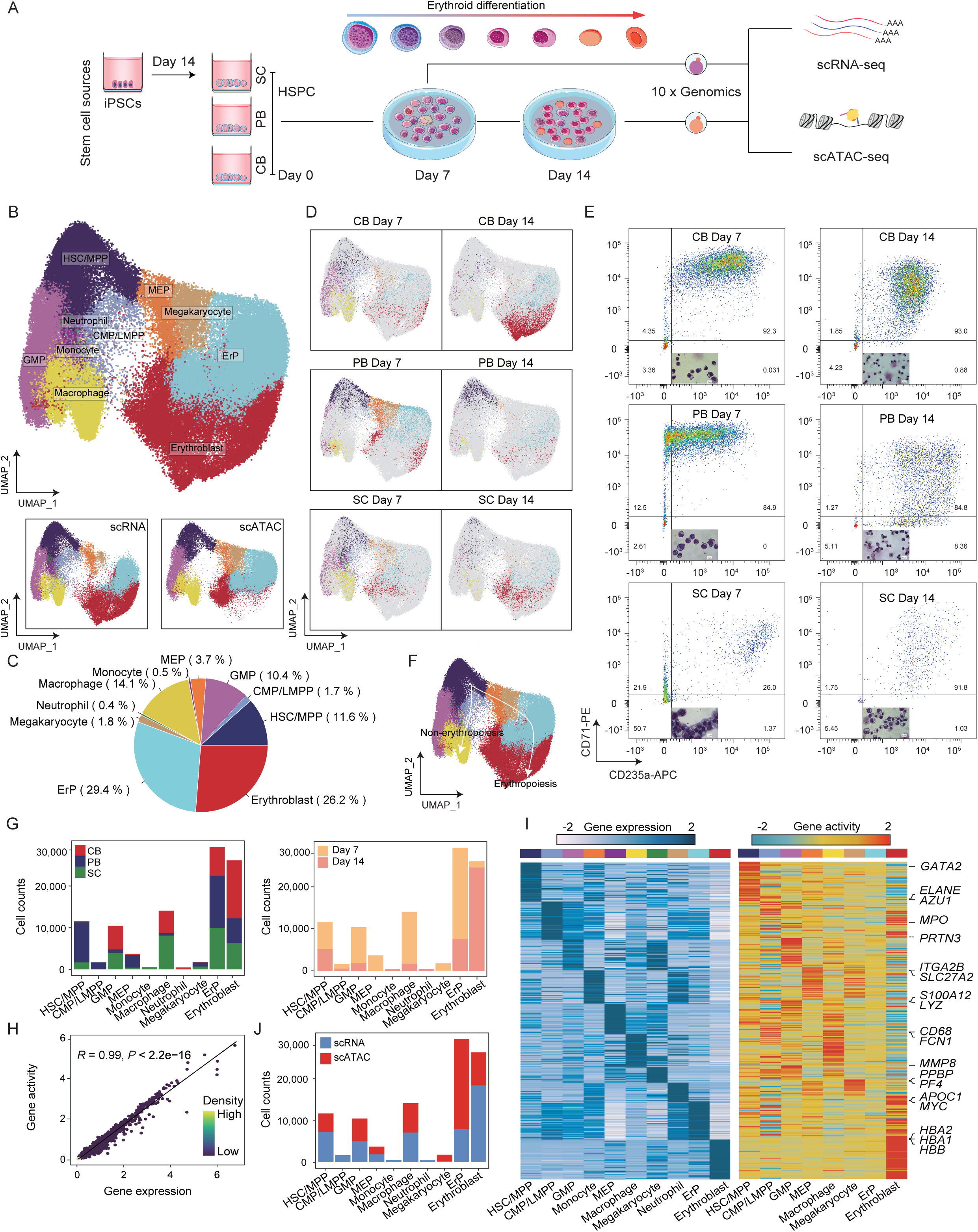
Single-cell chromatin accessibility and transcriptome maps of *ex vivo* erythropoiesis systems (see Figure S1 and Figure S2) A. Schematic illustration of the overall strategy of this study. B. Uniform manifold approximation and production (UMAP) plots of cell annotation results of scRNA-seq and scATAC-seq data. HSC/MPP, hematopoietic stem cells and multipotent progenitors; CMP/LMPP, common myeloid progenitors and lymphoid-primed multipotent progenitors; GMP, granulocyte-macrophage progenitors; MEP, megakaryocyte-erythroid progenitors; ErP, Erythroid progenitors. C. Pie chart showing the percentage of all cell types in Figure 1B. D. UMAP plots showing the cell distribution of different *ex vivo* erythropoiesis systems at two differentiation states. E. Flow cytometry results and morphology of erythroid cells on days 7 and 14 during the erythroid differentiation of CB-, PB-, and SC-derived systems. Scale bar = 10 μm. F. UMAP plot showing developmental pseudo-time trajectory from HSC/MPP to macrophages (non-erythropoiesis) and erythropoiesis (erythropoiesis). G. Bar plot showing the number of cell types in three *ex vivo* erythropoiesis systems (left) and at two differentiation time points (right). H. The correlation between the chromatin accessibility and expression of genes was analyzed based on the scRNA-seq and scATAC-seq data. The Pearson correlation was significant (*R* = 0.99, *P* < 2.2 × 10^−16^). I. Heatmap showing gene expression (left) and gene activity (right) of marker genes for different cell types. J. Bar plot showing the number of cell types measured by different single-cell sequencing methods.

To elucidate the production of myeloid cells within the *ex vivo* erythroid differentiation systems, we conducted a pseudo-time trajectory analysis and observed two main differentiation directions (erythropoiesis and non-erythropoiesis; Figure 1F, Additional file 1: Figure S2A, B) of HSC/MPP that highly expressed bilineage transcription factors (TF) including *GATA2* and *RUNX1* (Additional file 1: Figure S2C). Macrophages were mostly produced in the non-erythropoiesis direction (Figure 1C, Additional file 1: Figure S2A), with those accompanying RBCs generation during erythropoietin (EPO) induction of the *ex vivo* erythropoiesis.[5, 13] Along the direction of erythropoiesis, we observed increased expression of erythroid-specific genes (*GYPA* and *ALAS2*), while the expression of macrophage marker genes (*CD163* and *CD68*) was decreased (Additional file 1: Figure S2C). Likewise, along the direction of erythropoiesis, the accessibility of erythroid-specific TF motifs (e.g., *KLF1* and *NFE2*) increased, while that of specific transcription factor motifs (e.g., *GATA2* and *RUNX1*) associated with myeloid differentiation decreased (Additional file 1: Figure S2D).

To further investigate the origins of myeloid cells during *ex vivo* erythropoiesis, we analyzed the proportion of different cell types generated from *ex vivo* erythropoiesis systems. We observed that iPSC- and CB-derived erythropoiesis systems produced more myeloid cells (GMP and macrophages) compared to PB-derived system (Figure 1G), indicating a tendency toward myeloid lineage differentiation. However, during ex vivo erythroid differentiation from days 7 to 14, we noted a significant increase in erythroblast numbers and concomitant decrease in myeloid cell lineages (Figure 1G).

Furthermore, ErP were predominantly sampled on day 7, with a greater proportion of ErP from the CB-derived erythropoiesis system differentiating into erythroblasts compared to those from PB- and iPSC-derived systems, suggesting that the CB-derived system has the highest efficiency in inducing erythroid differentiation. Notably, the PB-derived erythropoiesis system exhibited the highest proportions of HSC/MPP and CMP/LMPP, which remained relatively stable throughout the erythroid differentiation process (Figure 1G).

We performed an integrative analysis of single-cell RNA sequencing and chromatin accessibility datasets. This analysis revealed significant concordance between the transcriptome and the chromatin accessibility profiles (Figure 1H), and robust expression and corresponding activity of marker genes across different cell types in the *ex vivo* erythropoiesis system (Figure 1I). This underscores the consistency between transcriptomic profiles and chromatin accessibility levels across cell types, implying the coordinated regulatory mechanisms governing *ex vivo* erythropoiesis. However, the identified erythroblasts were mostly derived from single-cell RNA sequencing (scRNA-seq) data, while the ErP were primarily derived from single-cell ATAC sequencing (scATAC-seq) data (Figure 1J). scATAC-seq captures chromatin accessibility using Tn5 transposase, which can be limited to capturing chromatin states in terminally differentiated erythroid cells.

### Trajectory analysis unveils myeloid differentiation bias in *ex vivo* erythropoiesis

To explore the potential mechanism behind myeloid production during *ex vivo* erythropoiesis, we establish early differentiation trajectories of *ex vivo* erythropoiesis systems by using HSPC (HSC, CMP/LMPP, GMP, and MEP) (Figure 2A). We divided the HSPC into three cell states. State 1 cells were located at the beginning of the pseudo- time trajectory (Additional file 1: Figure S3A), most of which were HSC/MPP from the PB-derived system (Figure 2B). Highly expressed genes in this state, including GATA2[14] and RHEX,[15] were enriched in erythroid differentiation, indicating the erythroid differentiation potential of HSC under EPO induction (Additional file 1: Figure S3B and S3C). State 2 cells with MEP mainly came from the PB-derived system, while state 3 cells with GMP and CMP/LMPP from the CB- and iPSC-derived systems, implying that HSC in CB and iPSC systems have a strong tendency for myeloid differentiation under EPO induction (Figure 2A and 2B). Consistently, we observed a higher enrichment of glutamine metabolism-related pathways (Figure 2C) and chromatin openness and expression of glutamine metabolism-related genes in the PB-derived system than those in CB- and iPSC- derived from state 1 cells (Figure 2D). Previous studies have revealed that the tendency of HSPC under EPO signaling is transferred to the myeloid cell fate while inhibiting glutamine metabolism, [12] implying that diminished glutamine metabolic activity may underlie the tendency of HSC in CB and iPSC systems to differentiate along the myeloid lineage, and iPSC and CB systems may enhance erythroid differentiation by activation of glutamine metabolism.

**Figure 2.**
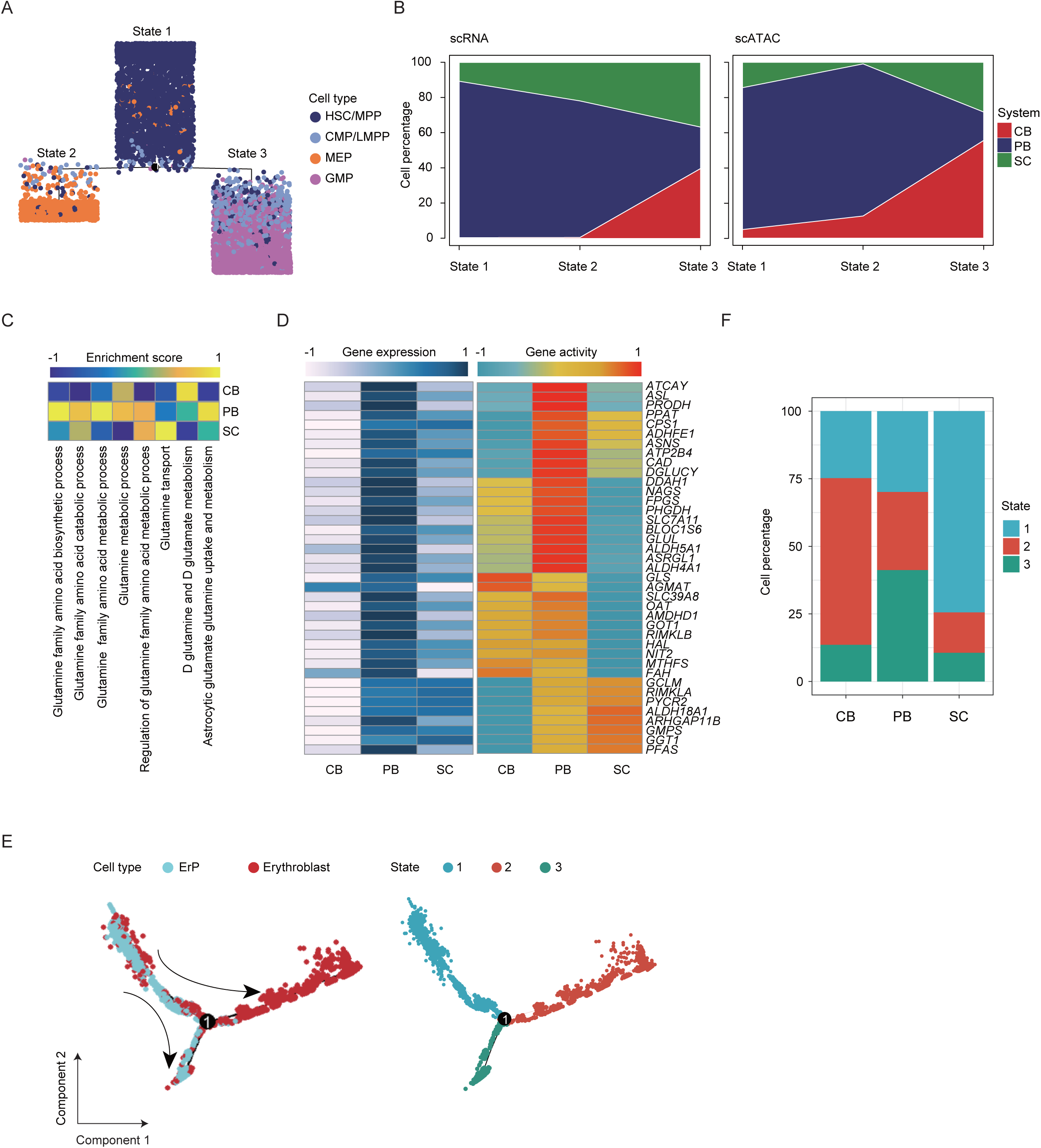
Myeloid differentiation tendency during *ex vivo* erythropoiesis (see Figure S3 and Figure S4) A. Distribution of hematopoietic stem and progenitor cells in the pseudo-time trajectory using Monocle 2. B. Proportion of hematopoietic stem and progenitor cells derived from the three *ex vivo* erythropoiesis systems in different states using scATAC-seq and scRNA-seq data respectively. C. Heatmap showing GSVA results of glutamine metabolism-related pathways in state 1 cells from different *ex vivo* erythropoiesis systems. D. Heatmap showing the activity scores and expression levels of glutamine metabolism-related genes in state 1 cells derived from different *ex vivo* erythropoiesis systems. E. Distribution of erythroid cells on a pseudo-time trajectory (Left) and distribution of three states on a pseudo-time trajectory (Right). F. The proportion of three states in Figure 2E from different *ex vivo* erythropoiesis systems.

We also plotted the differentiation trajectory of the erythroid lineage using ErP and erythroblasts and divided erythroid-related cells into three states using pseudo-time trajectory analysis (Figure 2E, Additional file 1: Figure S4A). The starting point of the differentiation trajectory was ErP, and two branches were formed over differentiation: erythroid cells on the main branch highly expressed *HBA1*, and erythroid cells on the secondary branch highly expressed *CD63* (Additional file 1: Figure S4B).[16] The analysis of the differentially expressed genes between the two branches (states 2 and 3) revealed that the secondary branch cells highly expressed myeloid lineage-related genes involved in neutrophil activation and leukocyte chemotaxis, while the main-branched erythroid cells highly expressed oxygen transport and mitochondrial autophagy-related genes (Additional file 1: Figure S4C and S4D). State 3 cells (most ErP) of the secondary branch activated specific TFs crucial for myeloid differentiation (CEBPA, CEBPB, SPIC). State 2 cells (most erythroid cells) were mostly derived from the CB-derived system, which specifically activated the GATA family TFs. State 1 cells (most CD63^+^TFRC^+^ ErP) in the pre-branch of differentiation were largely derived from the iPSC-derived system (Figure 2F, Additional file 1: Figure S4E). Collectively, our results revealed myeloid differentiation tendency of *ex vivo* erythropoiesis systems under the induction of EPO during *ex vivo* erythropoiesis.

### Lysosomal activity in stagnant HSC may impede *ex vivo* erythropoiesis

Residual HSC during *ex vivo* erythropoiesis system deteriorate erythrocyte regeneration efficiency. To characterized stalled HSC in *ex vivo* erythropoiesis systems, we compared the marker genes of HSC derived from *ex vivo* erythropoiesis systems and *in vivo* sources (BMMC and PBMC). Consistent with resting state of HSC *in vivo* sources, we found that the enhanced lysosomal activity was observed in HSC derived from the *ex vivo* PB-derived system, while the marker genes of HSC derived from BMMC and PBMC have a tendency to differentiate into lymphoid cells (Additional file 1: Figure S5A, S5B and S5C). In line with the enrichment of lysosome activity in the PB system, the chromatin accessibility and expression of lysosomal genes in PB system-derived state 1 cells derived from HSPC were higher than those in CB and iPSC system (Additional file 1: Figure S5D). Previous studies have shown that lysosomal pathway activation is crucial for maintaining the resting state of long-term HSC.[17] Lysosomes eliminate transferrin receptors on the surface of HSC, limiting their differentiation into the erythroid lineage[18] and causing them to enter the resting state. This finding suggests that the inactivation of lysosomal activity in the PB-derived system may increase the efficiency of erythroid differentiation.

### Erythroid atlas analysis reveals chromatin accessibility and gene expression coordination during *ex vivo* erythropoiesis

To investigate the coordination between chromatin accessibility and gene expression during ex vivo erythropoiesis, we conducted an in-depth erythroid atlas analysis. We employed a novel dimensionality reduction method, the potential of heat-diffusion for affinity-based transition embedding (PHATE),[19] for dimensionality reduction of erythroid cells, which can accurately reflect the cell differentiation trajectory while reducing the data dimension. The identified erythroid cells revealed a differentiation trajectory from BFU-E to CFU-E, followed by Proerythroblast (Pro-E), Basophilic erythroblast (Baso-E), Polychromatophilic erythroblast (Poly-E), and culminating in Erythroblast (Ortho-E) [Figure 3A, Additional file 3:Table S2].[8, 20, 21] Among erythroid cells, most ErP annotated by SCINA were defined as BFU-E or CFU-E, whereas most erythroblasts were defined as Poly-E and Ortho-E (Additional file 1: Figure S6A). Notably, more ErP were observed in the iPSC-derived system, while more erythroblasts were found in the CB-derived system (Figure 3B). The number of genes expressed sharply decreased during terminal erythroid differentiation (Figure 3C). Correspondingly, the average chromatin accessibility peaks measured during terminal erythroid differentiation also decreased (Figure 3D). The gene expression levels of different types of globin (embryonic, fetal, and adult) indicated the developmental stages during erythropoiesis. In contrast to embryonic and fetal globin, the expression of adult-type globin increased during *ex vivo* erythropoiesis, as evidenced by increased chromatin openness on β-globin during erythropoiesis (Figure 3E, Additional file 1: Figure S6B). Correlation analysis between the transcriptome and chromatin accessibility at developmental stages during erythroid differentiation revealed greater similarity in gene expression among these cell types and weaker similarity in chromatin accessibility (Figure 3F). This suggests a greater regulatory amplitude of chromatin accessibility during *ex vivo* erythroid differentiation.

**Figure 3.**
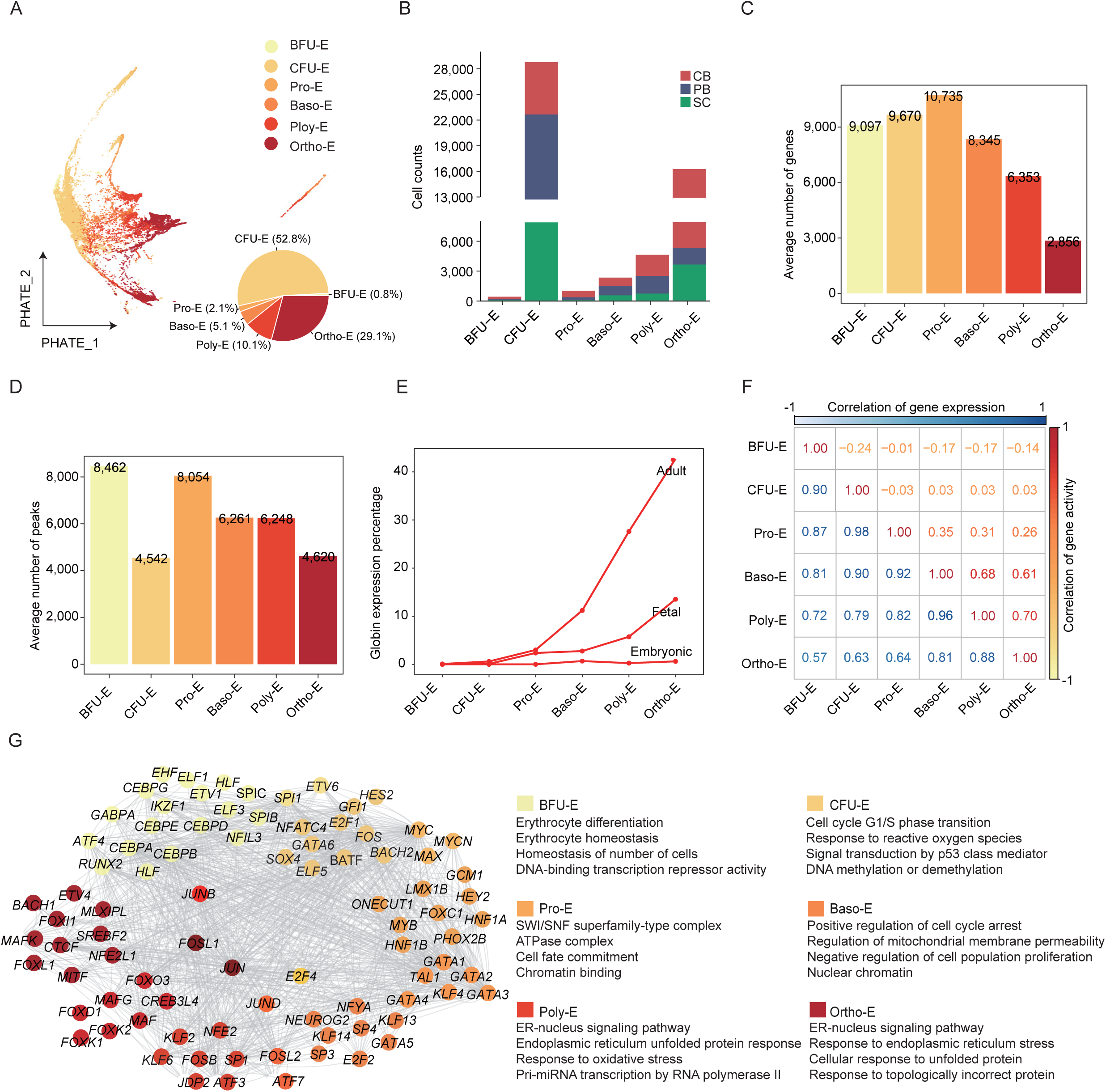
Atlas of erythroid cells of *ex vivo* erythropoiesis systems (See Figure S6) A. Dimensionality reduction plot of the erythroid cell subpopulations visualized using PHATE. Pie chart showing the percentage of erythroid cell subpopulations. B. Bar plot showing the number of erythroid cell subpopulations from three erythropoiesis systems. C. Bar plot showing the average number of genes in each erythroid cell subpopulation. D. Bar plot showing the average number of peaks in each erythroid cell subpopulation. E. Mean expression levels of adult-, fetal-, and embryo-type globin in erythroid cell subpopulations. F. Correlation between gene expression and gene activity score among erythroid cell subpopulations. G. Regulatory network and GO-enriched pathways of TFs in each erythroid cell subpopulations.

To further elucidate the role of chromatin accessibility during *ex vivo* erythropoiesis, we identified the characteristic peaks of chromatin accessibility across different developmental stages. Additionally, using ChromVAR, we identified motif-enriched chromatin accessibility variability associated with each erythroid cell type and ascertained the enrichment of erythroid-specific TFs at each developmental stage (Additional file 1: Figure S6C). More TF motifs were enriched in ErP (especially CFU-E) than in terminal erythroblasts, indicating that more TFs were required to drive erythroid differentiation (Additional file 1: Figure S6D). Next, we used the TFs (top 20) specifically highly activated at each developmental stage, and mapped the TFs regulatory network corresponding to each developmental stage (Figure 3G, Additional file 3: Table S2). The TFs (*MYC*, *MAX*, *E2F1*) specifically activated in CFU-E are related to cell cycle transition and response to oxygen, whereas the TFs (*E2F2*, *JUND*, *NYFA*) specifically activated in Baso-E potentially regulated cell cycle arrest and nuclear chromatin. The highly activated TFs regulatory network of erythroid cells revealed that chromatin accessibility during erythroid differentiation was stage-specific and significantly changed.

### Cell cycle arrest and hypoxia signaling deficiency impair erythroid differentiation of ErP during *ex vivo* erythropoiesis

In our dataset, 456 burst-forming unit erythroid (BFU-E) and 29,314 colony-forming unit erythroid (CFU-E) were identified. These cells mainly originated from samples on day 7, and Poly-E and Ortho-E cells in the late stage of terminal differentiation mainly originated from samples on day 14 (Figure 4A). However, a population of CFU-E and BFU-E were still in the sample on day 14, mainly from the iPSC-derived erythropoiesis system, followed by the PB-derived system; the CB system had the least residual ErP (Figure 4B). To explore the underlying restraints that prevented ErP from differentiating downward, we characterized ErP across different *ex vivo* erythropoiesis systems and between the two differentiation states at the levels of chromatin openness and gene expression (Figure 4C, Additional file 1: Figure S7A). Compared with the PB- and iPSC-derived systems, ErP derived from the CB-derived system exhibited a significant enrichment of highly expressed and active genes in the cell cycle checkpoint pathway (Additional file 1: Figure S7B and S7C). Moreover, genes with higher expression and chromatin accessibility in ErP on day 7 compared to day 14 were significantly enriched in pathways associated with hypoxia-inducible factor 1 (HIF-1) and cell cycle regulation (Figure 4D and 4E). This suggests that early erythroid progenitors may rely on these pathways to maintain their proliferative and differentiation potential during ex vivo erythropoiesis.

**Figure 4.**
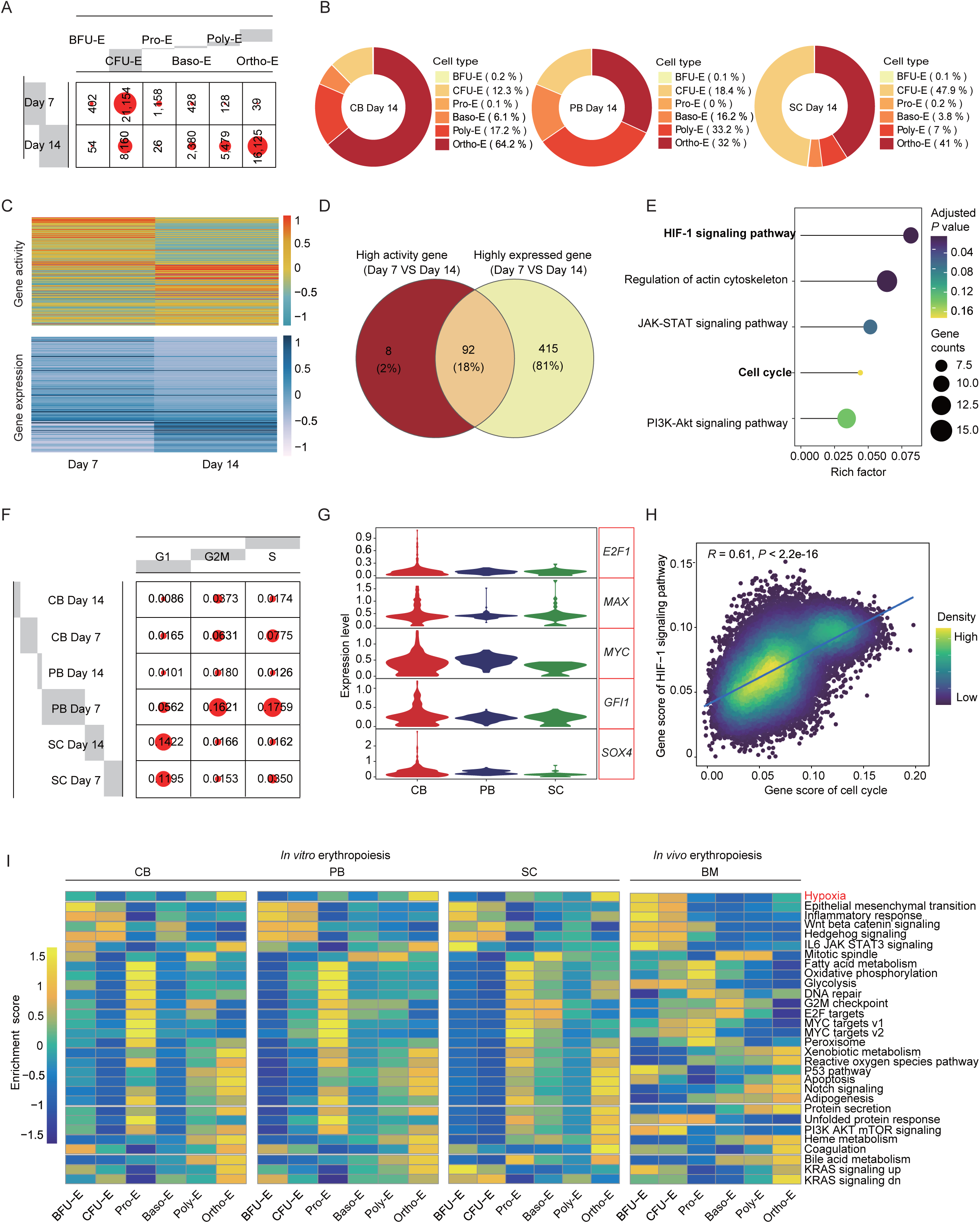
Analysis of impaired ErP differentiation potential in *ex vivo* erythropoiesis systems (see Figure S7) A. Cell number of each erythroid cell subpopulation on days 7 and 14. B. Proportion of each erythroid cell type on day 14 from different *ex vivo* erythropoiesis systems. C. Heatmap showing gene activity scores and expression levels of ErP (BFU-E and CFU-E) on days 7 and 14. D. Venn diagram showing the overlap between highly active and highly expressed genes on day 7 compared with that of day 14 in ErP. E. GO-enriched terms of genes intersected in Figure 3D. F. Proportion of ErP at different cell cycle phases for each sample. G. Violin Diagram showing the expression levels of *E2F1*, *MYC*, and other TFs-encoding genes that regulate cell cycle of ErP in different erythropoiesis systems. H. Correlation between the cell cycle and HIF-1 signaling pathway. The Pearson correlation was significant (*R* = 0.61, *P* < 2.2 × 10^−16^). I. Heatmap showing the GSVA score in each erythroid cell subpopulation from different *ex vivo* erythropoiesis systems compared with that of healthy BM.

Further cell cycle analysis revealed that approximately 8% of CB system-derived ErP on day 7 were in the S phase, whereas ErP system derived from the iPSC system on the same day exhibited 3.5% of ErP in the S phase, with a higher proportion in the G1 phase. Furthermore, most ErP on day 14 were arrested in the G1 phase (Figure 4F), suggesting that impaired differentiation of ErP in the iPSC-derived system is likely attributable to cell cycle dysfunction. Subsequent investigations demonstrated that TFs (e.g., E2F1, MAX, MYC) crucial for cell cycle regulation are highly expressed in ErP derived from the CB system (Figure 4G). Moreover, the iPSC system appeared to have deficiencies in both cell cycle regulation and HIF-1 signaling (Figure 4F, Additional file 1: Figure S7D), with a positive correlation between these two pathways (Figure 4H).

We subsequently analyzed the enriched pathways in ErP from the *ex vivo* erythropoiesis systems comparing with those in BM. The pathway profiling of BFU-E and CFU-E were similar across three *ex vivo* erythropoiesis systems but markedly differed from those in the BM. A pronounced divergence in the response to oxygen was observed between ErP derived from *in vitro* and *in vivo*. Specifically, BM-derived ErP exhibited significant enrichment in hypoxia signaling (Figure 4I). Hypoxia signaling in CB-derived BFU-E was more robust than that in cells derived from PB- and iPSC-derived erythropoiesis systems, suggesting a deficiency in HIF-1 signaling within PB- and iPSC-derived ErP (Figure 4I). Collectively, the impaired hypoxia signaling observed in *ex vivo* erythropoiesis systems, particularly in iPSC-derived system, may present a significant constraint on the differentiation of ErP during *ex vivo* erythropoiesis.

### Chromatin accessibility-associated enhancers and their regulatory role in *ex vivo* erythropoiesis

To further explore the regulatory role of chromatin accessibility during *ex vivo* erythropoiesis, we performed a linkage analysis of chromatin accessibility peaks with all genes expressed in erythroid cells (Figure 5A, Additional file 4: Table S3) and identified a total of 44,062 linkage pairs (z-score > 0, p < 0.05). K-means clustering divided the linkage pair into six clusters, in which the enhancer activity in cluster 2 significantly decreased during *ex vivo* erythropoiesis, and the enhancer activity in cluster 6 significantly increased (Figure 5B, Additional file 4: Table S3). Consistent with the current findings (Figure 3G), TFs, including HIF1α, MYC, and MAX (Figure 5C, Additional file 4: Table S3), were enriched in the peaks in cluster 2 that were responsive to erythroid differentiation and erythrocyte homeostasis (Additional file 1: Figure S8A-C),[22–24] implying that the enhancers in BFU-E can determine the erythropoiesis pathway. The corresponding chromatin accessibility peaks in cluster 6 enhancers that could be enriched by TFs, including KLF1, KLF6 and FOXO3, were specifically opened (Figure 5D, Additional file 4: Table S3), and the potentially regulated genes were enriched for oxygen transport, iron ion transport, and autophagy,[25, 26] implying that the enhancers in cluster 6 are pivotal for *ex vivo* erythroid maturation (Additional file 1:Figure S8A-C).

**Figure 5.**
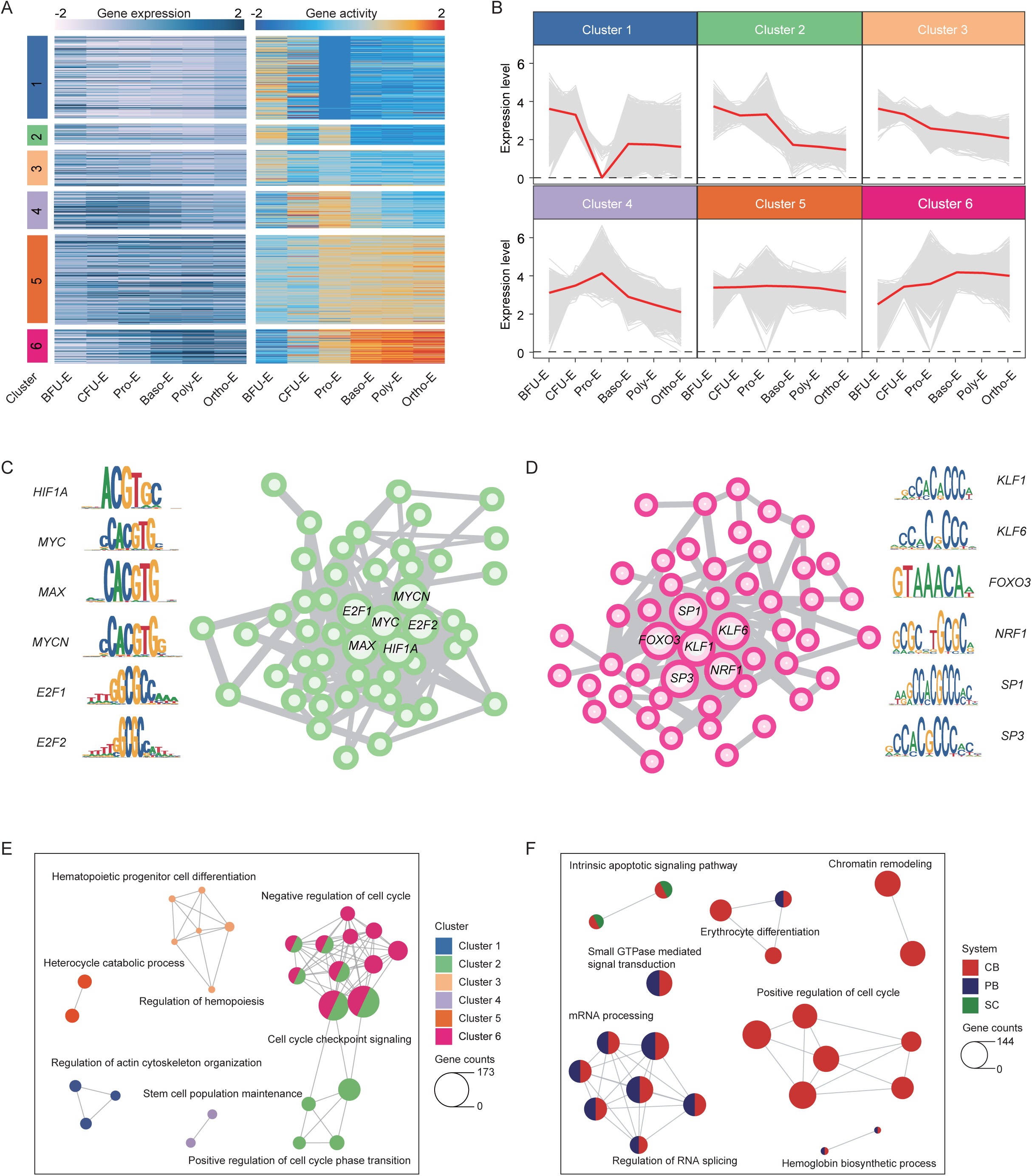
Chromatin accessibility-associated enhancers and their regulatory role in ex vivo erythropoiesis (see Figure S8) A. Heatmap showing clustering comparison based on the abundance of chromatin accessibility peaks and expression levels of the genes in erythroid cells. B. Line plots showing the expression dynamics of genes potentially regulated by enhancers during *ex vivo* erythropoiesis. C. Regulatory network of transcription factors enriched on peaks in cluster 2. D. Regulatory network of transcription factors enriched on peaks in cluster 6. E. The network showing the functional annotation of genes corresponding to the intersecting peaks between each link cluster’s peaks and the system’s marker peaks, as depicted in Figure S8D. The colors are filled according to the link clusters. F. The network showing the functional annotation of the intersecting peaks, with colors filled according to the different systems.

To elucidate the system origins of these enhancer peaks, we performed intersection analyses between system-specific marker peaks and enhancer peaks from each erythropoiesis system, and defined overlapping peaks as system-specific enhancer peaks. We identified a greater abundance of enhancer peaks in the CB-derived erythropoiesis system relative to the PB- and iPSC-derived systems (Additional file 1: Figure S8D). Interestingly, system-specific enhancer peaks tended to be proximal to the promoter regions, whereas non-system-specific enhancer peaks were predominantly distal to these regions (Additional file 1: Figure S8E and S8F), suggesting that system-specific enhancer peaks possess robust gene-regulatory capabilities. Functional annotation network analysis indicated that the genes regulated by system-specific enhancer peaks in cluster 2 were closely associated with cell cycle regulation. Moreover, cell cycle-related pathways were more enriched in umbilical CB systems than they were in PB- and iPSC-derived erythropoiesis systems (Figure 5E and 5F).

### Enhancer-gene interactive regulation of erythroid maturation

Our study found that erythroid cells derived from the iPSC-derived erythropoiesis system exhibit a low proportion of total globin expression, encompassing the adult, fetal, and embryonic types. Conversely, erythroid cells from the PB exclusively express adult-type globin. Erythroid cells from the CB system also express adult-type globin, retain fetal-type globin expression, and lack embryonic globin (Figure 6A). Since the globin gene was the most highly expressed in Ortho-E, we used globin expression-related genes recorded in the database to reduce the dimensionality of Ortho-E and obtained four types of terminal erythroblast populations. The ε-Ortho-E subpopulation, characterized by elevated expression of the *HBZ* gene, was uniquely identified within the iPSC-derived erythropoiesis system. Comparatively, γ-Ortho-E, marked by elevated levels of *HBG1* expression, was predominantly sourced from the CB-derived erythropoiesis system. Additionally, the β-Ortho-E subpopulation, distinguished by robust *HBB* expression, was predominantly represented in both the PB- and CB-derived erythropoiesis systems. The Reti-E subpopulation, which expresses both *HBG1* and *HBB*, was derived exclusively from scRNA-seq data (Figure 6B and 6C). We further analyzed the chromatin accessibility peaks and expression patterns of all β-globin genes, including *HBE1*, *HBG1/HBG2*, *HBD*, and *HBB*, along with the identification of their potential enhancer regions within distinct Ortho-E subpopulations across the β-globin locus (Figure 6D and 6E). For β-Ortho-E, we identified two predominant enhancer peaks within the genomic region of the globin locus. These enhancer regions were highly correlated with the TFs including TAL1 and RORA (Figure 6F). These findings suggest a putatively interactive network for β-globin expression, where functional TFs (e.g., RORA, TAL1) are implicated through their binding to specific motifs within *HBB* enhancers.

**Figure 6.**
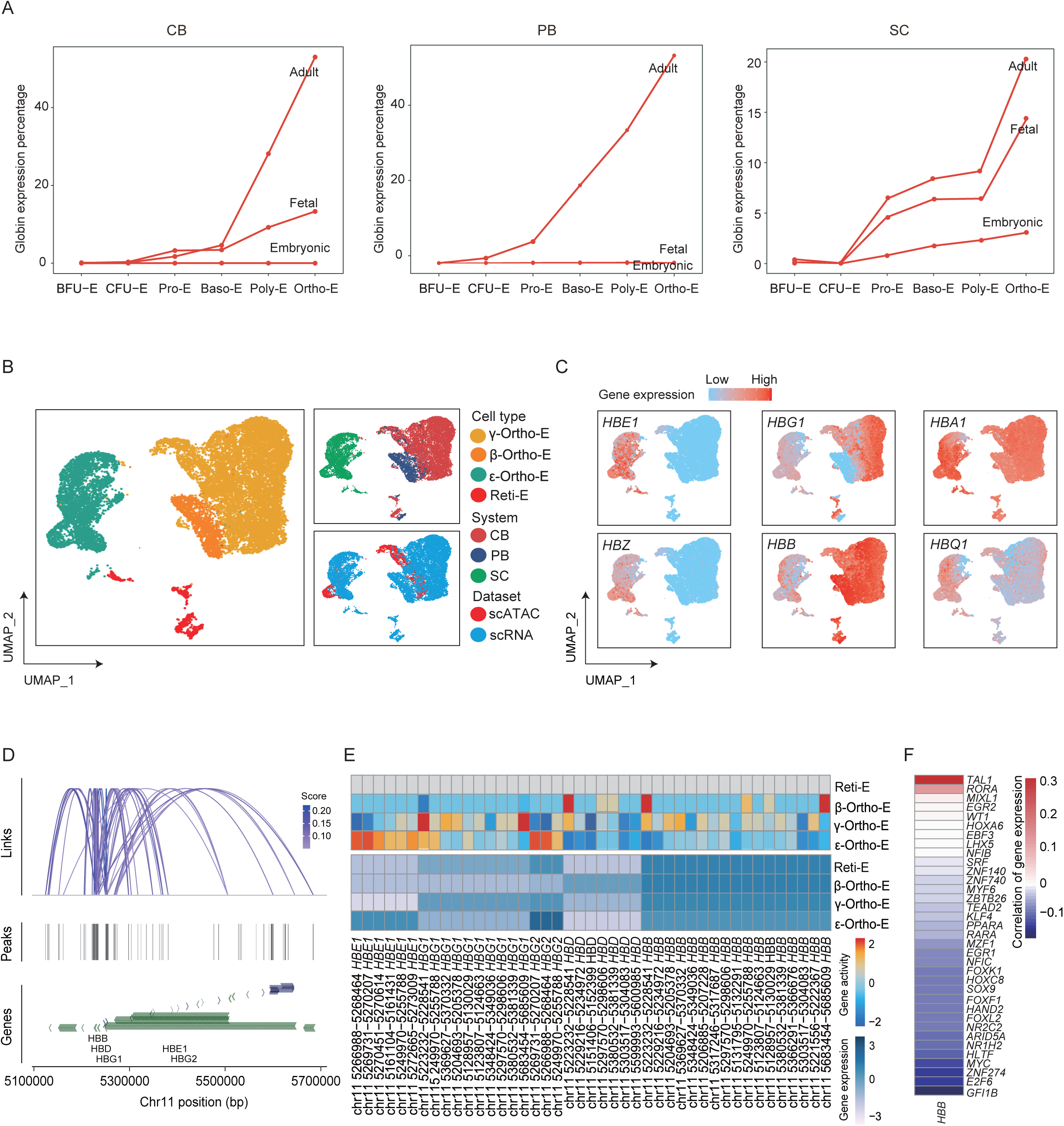
Enhancer-gene interactive regulation of globin gene expression. A. Line plots showing the mean expression levels of adult-, fetal-, and embryo-type globin-encoding genes in erythroid cell subpopulations of different erythropoiesis systems. B. UMAP plots of Ortho-E and reticulocytes across different erythropoiesis systems and datasets. C. UMAP plots of Ortho-E and reticulocytes revealing the expression patterns of globin genes (*HBE1*, *HBZ*, *HBG1*, *HBB*, *HBA1*, and *HBQ1*). D. Chromatin accessibility peaks on β-globin coding genes, including *HBB*, *HBD*, *HBG1*, *HBE1*, and *HBG2*. Normalized scATAC-seq sequencing tracks of erythroid cell subpopulations at gene loci. The upper panel is the score of co-accessible links. E. Heatmap showing the expression level of β-globin-encoding genes (*HBG1*, *HBG2*, *HBD*, and *HBB*) and their potential enhancer peaks in Ortho-E and reticulocytes. Upper: potential enhancer peaks; lower: gene expression levels. F. Heatmap showing the expression correlation of enriched TFs on *HBB* potential enhancers with *HBB*.

Erythroid enucleation remains a significant challenge in erythrocyte regeneration.[2] Our analysis of terminal erythroid cells revealed that Ortho-E and reticulocytes (Reti-E) predominantly originate from the CB-derived system, indicating that these cells from the CB system possess a more robust enucleation capacity compared with those derived from PB- or iPSC-derived erythropoiesis systems. We explored chromatin accessibility in Ortho-E to uncover the potential regulatory mechanisms underlying erythroid enucleation. We compared chromatin accessibility and gene expression in Ortho-E across *ex vivo* erythropoiesis systems and found a significant association between high chromatin openness and gene expression in the CB-derived system, particularly for genes involved in chromosome condensation and autophagy, a process critical for erythrocyte enucleation during terminal erythroid differentiation (Additional file 1: Figure S9A-C). Previous studies have shown that autophagy plays a key role in the elimination and enucleation of organelles such as mitochondria in the process of terminal erythroid differentiation, in which FOXO3 is an important regulator of autophagy[27] and is highly expressed with increased chromatin accessibility in the CB-derived erythropoiesis system (Additional file 1: Figure S9D). Certain enhancers with increased chromatin accessibility are known to transcribe enhancer RNAs (eRNAs), a form of long non-coding RNA (lncRNA) ranging from 0.5–5 kb in length. Analysis of potential enhancer-proximal genes across erythroid cell stages revealed a significant proportion of lncRNAs (Additional file 1: Figure S8B and S8C). We identified marker lncRNAs in Ortho-E, including MALAT1, MIR22HG, and MIAT, that are specifically associated with an enhancer from cluster 6 and regulate FOXO3 (Additional file 1: Figure S9D, S9E-G; Additional file 3: Tables S2; Additional file 4: Table S3). To identify the downstream target genes regulated by the TFs in Ortho-E, we utilized the SCENIC algorithm to infer and screen for target genes with elevated expression levels within Ortho-E. These genes were correlated with lncRNAs, leading to the construction of an enhanced RNA (eRNA)– mRNA regulatory network implicated in erythrocyte enucleation (Additional file 1: Figure S9H). TMCC2 is integral to terminal erythroid differentiation; its deficiency significantly reduces enucleated erythroid cells.[28] These results imply that the eRNA- TFs-mRNA interacting network in ortho-E may regulate erythroid enucleation, particularly for iPSC- or PB-derived erythropoiesis systems.

### Differential cell-cell interactions between *ex vivo* and *in vivo* erythropoiesis systems

We explored cell-cell interactions to gain deeper insights into the underlying constraints for *ex vivo* erythropoiesis. We compared the cell–cell interaction profiles of the microenvironment of the erythropoiesis system between *in vivo* and *in vitro*, and found a significantly enhanced role of erythroid cells during *ex vivo* erythropoiesis (Figure 7A and 7B, Figure S10A). In the BM, fewer molecules interact between cells, while the intensity of these interactions was high; conversely, in the *in vitro* erythropoiesis system, a greater number of molecules interact between cells, but with a lower intensity, possibly due to the low expression level in the *in vitro* microenvironment (Additional file 1: Figure S10A and S10B).

**Figure 7.**
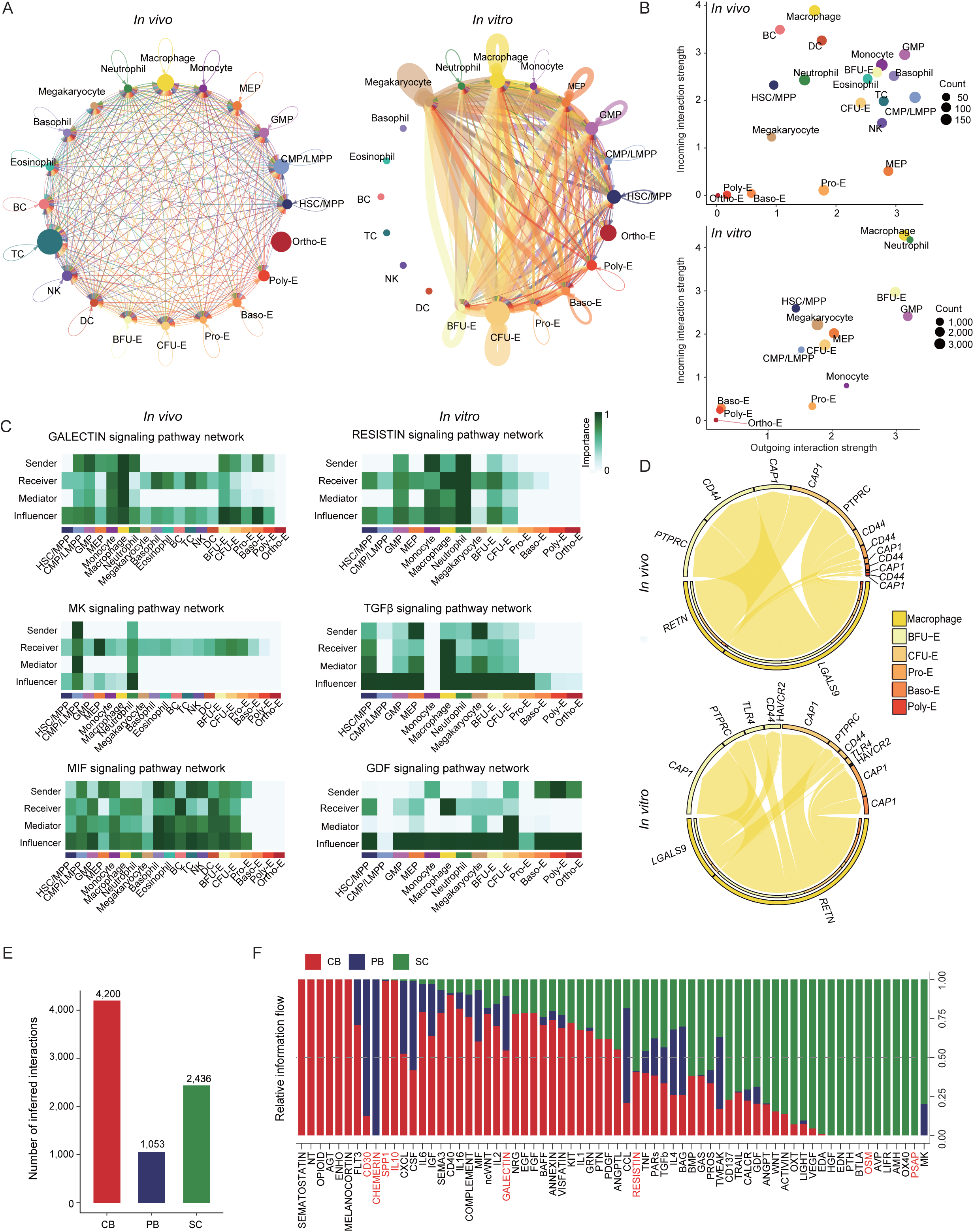
Cell–cell interaction map of erythropoiesis systems *in vivo* and *in vitro* (see Figure S10) A. Circle plots showing differential cell–cell communication networks *in vivo* and *in vitro.* The width of the edges represents the relative number of interactions or interaction strength. The scRNA-seq data were used. *In vitro*, the erythropoiesis systems (days 7 and 14) derived from CB, PB, and iPSC; *In vivo*, BM hematopoietic cells from healthy individuals (GSE150774 and GSE139369). B. Scatter plots showing dominant senders (sources) and receivers (targets) in a 2D space. The x- and y-axes represent the total outgoing or incoming communication probabilities associated with each cell group (*in vivo* and *in vitro*). Dot size is proportional to the number of inferred links (both outgoing and incoming) associated with each cell type. Dotted colors indicate different cell types. C. Different roles of each cell type in specific pathways. GALECTIN, MK, and MIF signaling pathways *in vivo* and RESISTIN, TGFβ, and DGF signaling pathways *in vitro*. D. Cell–cell communications between erythroid cells and macrophages *in vivo* and *in vitro*. E. Number of inferred communication links between different erythropoiesis systems. F. Ranking signaling networks based on information flow. Significant signaling pathways were ranked based on differences in the overall information flow within the inferred networks between the different erythropoiesis systems.

We profiled the signaling pathways involved in cell–cell interactions for in vitro with in vivo microenvironments and found several key differential pathways (Additional file 1: Figure S10C). We further compared the cellular and molecular contributions of the in vivo cell-specific interaction pathways, including GALECTIN, MK, and MIF, and *in vitro* cell-specific interaction pathways, including RESISTIN, TGFβ, and GDF (Figure 7C, Additional file 1: Figure S10C and S10D; Additional file 5:Table S4), and found that macrophages communicate with erythroid cells mainly through LGALS9-CD44 and LGALS9-CD45 *in vivo*, while communication with erythroid cells was mainly through RETN-CAP1 *in vitro* (Figure 7D). Macrophages are the most non-erythroid cell population in *ex vivo* erythropoiesis systems, and cell–cell contacts between macrophages and erythroid cells facilitate erythroid maturation.[29, 30] Galactoglutinin (GALECTIN) plays an important role in erythroblast islands,[31] whereas CAP1 inhibits the differentiation and enucleation of erythroid cells.[32] These results suggest that a good erythropoiesis microenvironment is not successfully established in *in vitro* systems.

We further compared cell–cell contacts across the three systems and revealed significant differences in the communication modes among three *ex vivo* erythropoiesis systems. The ligands secreted by cells *in vitro* primarily originated from CB- and iPSC-derived erythropoiesis system, which may be related to their strong tendency toward myeloid differentiation (Figure 7E). We observed the specific signals in distinct *ex vivo* erythropoiesis system: SPP1 and IL10 signals in CB-derived system, CHEMERIN and CD30 signals in PB-derived system, and PSAP and OSM signals in iPSC-derived system. We observed the highest enrichment of the GALECTIN pathway in CB-derived system but the weakest enrichment in iPSC-derived system; moreover, we observed highly conserved RESISTIN pathway enriched in iPSC- and CB-derived systems but not in PB-derived system (Figure 7F). These results suggest that modulating cell-cell communication signals in distinct *ex vivo* erythropoiesis system may help to build an effective microenvironment for erythrocyte regeneration.

## Discussion

Bioinformatics analysis of *ex vivo* human erythropoiesis from different sources offers insights into erythroid differentiation and elucidates intervention targets to enhance RBCs production.[2, 5, 6, 8, 11] While RBC regeneration from diverse sources yields suboptimal erythroid cells with variable maturation and efficiency,[33] few studies have decoded the discrepancies in *ex vivo* human erythropoiesis at single-cell multi-omics resolution or uncovered the underlying constraints of RBC regeneration.

Myeloid cells play an important role in erythroid differentiation,[34, 35] though the presence of myeloid cells might affect cell purity. Interestingly, in iPSC system, a significant proportion of HSC differentiate toward the myeloid lineage. iPSC differentiation systems have lower glutamine metabolism, suggesting a link between this shift toward the myeloid lineage and low glutamine metabolism that determines the erythroid fate of HSC. Erythroid differentiation of HSC requires the glutamine transporter protein, ASCT2, by facilitating glutamine uptake; blocking this pathway allows EPO-stimulated HSC to differentiate into the myeloid lineage.[36] In addition, the utilization of glutamine is necessary for erythroid maturation.[37] Glutamine supplementation could reduce myeloid cells in the *ex vivo* erythropoiesis systems (iPSC or CB), and may facilitate the maturation of erythroid cells.

Our study revealed that approximately 50% of ErP were stranded in the iPSC differentiation system even at the late stage (Day 14) during terminal erythroid differentiation. The other groups also revealed a comparable percentage of ErP in differentiation systems derived from CB, PB, and iPSC at the same time point.[23, 38, 39] ErP have been even identified from the terminal erythroid differentiation system on days 21–23 from CB- and iPSC-derived erythropoiesis systems,[5, 39] indicating that stranded ErP restrained erythrocyte production. In the current study, we found that most ErP were in the G1 phase, and TFs (*E2F1*, *MYC*) related to the cell cycle and cell proliferation were suppressed in the ErP of the iPSC system, implying cell cycle arrest in the iPSC-derived system, consistent with previous finding that the cell cycle arrest in ErP impaired erythroid proliferation and differentiation under pathologic state. [40] As such, supplement of antagonists for cell cycle arrest or regulation of signaling pathway controlling cell cycle arrest may be the strategies to improve *ex vivo* erythropoiesis by transforming stranded ErP into mature erythroid cells. Moreover, oxygen levels regulate cell proliferation depending on the cell type. We found that hypoxia signaling was closely associated with cell cycle arrest in ErP, implying that hypoxia could inhibit cell proliferation by provoking cell cycle arrest. [41] The hypoxia signaling in the iPSC-derived system were significantly suppressed compared with ErP from the other *in vitro* differentiation systems and BM. Previous experimental studies demonstrated that ErP achieved a remarkably high expansion efficiency,[39] along with the maturation of RBCs under hypoxic conditions.[23, 39] The activation of the hypoxia signaling may be beneficial for the expansion and differentiation of ErP in the iPSC-derived erythropoiesis system.

Chromatin accessibility has rarely been used to explore the mechanisms of restrained RBCs regeneration. In this study, we profiled the chromatin accessibility peaks along all beta-globin loci for Ortho-E classified based on globin expression levels, and found an obvious discrepancy of chromatin accessibility peaks existing in the genomic region of ε-, γ-, and β-globin among three *ex vivo* erythropoiesis systems. Ortho-E derived from the PB-derived system demonstrated exclusive *HBB* expression that was abolished in the iPSC system and compromised in the CB system, consistent with previous observation.[42] The enhancer and gene interactive regulation in the PB-derive system could be implemented in iPSC or CB-derived erythropoiesis systems to enhance adult globin expression. Strategies to augment chromatin accessibility at specific enhancer regions, for example, by enhancing histone acetylation [43, 44] or changing DNA modifications with small molecules,[45] maybe ameliorate adult hemoglobin expression in these differentiation systems.

Although the *ex vivo* culture attempts to mimic the *in vivo* conditions as closely as possible, it is possible to obtain a certain order of magnitude of red blood cells with typical morphology and certain functions, yet there remains a gap between the *ex vivo* - induced red blood cells and those naturally grown *in vivo*.[16] The *in vivo* erythropoiesis is extraordinarily complex,[46] involving intricate cell interactions and microenvironmental factors that are often overlooked in *ex vivo* cultures. Macrophages, for example, are integral to the production of red blood cells in our body.[47] Our data indicate that erythropoiesis *in vivo* is even more complex than previously understood, with significant differences in the molecular pathways of interaction between macrophages and erythroid cells *in vivo* versus *ex vivo*. The discrepancies in the interaction pathways of GALECTIN and RETN-CAP1 between *in vivo* and *ex vivo* differentiation systems reveal this complexity, indicating that erythropoiesis *in vivo* is far more intricate than currently understood. The forthcoming study will continue to explore the role of the microenvironment in erythropoiesis, both *in vivo* and *ex vivo*, to better simulate the *in vivo* conditions and develop a regenerative system that more closely mimics the natural BM environment, thereby enhancing regenerative outcomes.

## Conclusions

This study provides a comprehensive single-cell analysis of *ex vivo* human erythropoiesis from CB, PB, and iPSC sources, revealing key discrepancies and constraints for erythrocyte regeneration. The *ex vivo* erythropoiesis systems from different sources exhibit distinct myeloid differentiation tendencies, and these tendencies were associated with low glutamine activity in cord blood- and iPSC-derived erythropoiesis. Cell cycle arrest and hypoxia signaling deficiency may lead to the stranded ErP in the iPSC-derived system. We observed the coordinated regulation between chromatin accessibility and transcriptome and the pivotal role of chromatin accessibility and its associated enhancers in regulating *ex vivo* erythropoiesis. Cell–cell communications in the *ex vivo* erythropoiesis system were not as well established as those in the BM. These insights suggest that modulating glutamine metabolism, cell cycle regulation, and cell-cell communication signals may improve *ex vivo* erythropoiesis. Our findings have significant implications for regenerative medicine.

## Methods

### iPSC culture and derived erythropoiesis

iPSCs were cultured in vitronectin (Catalog No. A14700, Invitrogen, Carlsbad, CA, USA)-coated culture dishes using Essential 8 Medium (Catalog No. A1517001, Gibco, Carlsbad, CA). When the cells reached 70%–80% confluence (usually around 3–4 days), they were digested to suspension cells (SCs) using 0.5 mM EDTA (Catalog No. 15575020, Invitrogen) and passaged at a ratio of 1:4. EB culture was performed from days 0 to 11 as previously described [11, 48]. From days 11 to 14 of the EB culture, thrombopoietin (TPO) was replaced with 3 U/mL EPO (Catalog No. 100-64, PeproTech, Cranbury, NJ). On day 14, SCs were collected using a 70-μm cell strainer and analyzed using flow cytometry using anti-human CD34-PE (Catalog No. 12-0349-41, eBioscience) and anti-human CD45-FITC (Catalog No. 560976, BD). Subsequently, the SCs were centrifuged at 300 × *g* for 5 min to remove the supernatant and washed once with 2% FBS. For erythroid differentiation, SCs were cultured at 10^5^ cells/mL for 7 days (days 15–21 of the whole process) in SFEM II medium supplemented with 1% penicillin/streptomycin (Catalog No. 15140122, Life Technologies, Carlsbad, CA, USA), 10 ng/mL stem cell factor (SCF; catalog No. 300-07, PeproTech), 10 ng/mL IL-3 (Catalog No. 200-03, PeproTech), and 3 U/mL erythropoietin (EPO). Then, the cells were cultured at 4 × 10^5^ cells/mL for 4 additional days (days 22–25 of the whole process) in SFEM II supplemented with SCF and EPO, and cultured at 7.5 ×10^5^ cells/mL for 3 additional days (days 26–28 of the whole process) in SFEM II supplemented with EPO. Erythroid cells were differentiated by incubation at 37 ℃ and 5% CO_2_.

### HSPC isolation and derived erythropoiesis

To enrich HSPC from fresh cord blood and peripheral blood samples, we isolated CD34+ cells from human cord blood using an EasySep™ Human Cord Blood CD34 Positive Selection Kit II (Catalog No. 17896, STEMCELL Technologies, Vancouver, Canada), and from human peripheral blood using an EasySep™ Human CD34 Positive Selection Kit II (Catalog No. 17856, STEMCELL Technologies, Vancouver, Canada). A sample purity of > 95% was confirmed using flow cytometry. When the expansion phase was required, CD34+ cells were cultured in StemSpan SFEM II medium (STEMCELL Technologies) supplemented with 50 ng/mL stem cell factor, 50 ng/mL Flt3 ligand (PeproTech, Rocky Hill, NJ, USA), 50 ng/mL thrombopoietin (PeproTech), and 1% penicillin and streptomycin solution for 7 days prior to differentiation. CD34+ cell purity was > 95% after expansion, as determined using fluorescence-activated cell sorting. For erythroid differentiation, CD34+ cells were cultured at 10^5^ cells/mL for 7 days in SFEM II medium supplemented with 1% penicillin/streptomycin (Catalog No. 15140122, Life Technologies, Carlsbad, CA), 10 ng/mL stem cell factor (SCF; catalog No. 300-07, PeproTech), 10 ng/mL IL-3 (Catalog No. 200-03, PeproTech), and 3 U/mL erythropoietin (EPO). Cells were then cultured at 4 × 10^5^ cells/mL for 4 days in SFEM II supplemented with SCF and EPO, and cultured at 7.5 ×10^5^ cells/mL for 3 additional days in SFEM II supplemented with EPO. Erythroid cells were differentiated by incubation at 37 ℃ and 5% CO_2_. Cord blood samples from full-term newborns were obtained from the Beijing Cord Blood Bank (China). Human peripheral blood samples from healthy donors were obtained from the Chinese PLA General Hospital in Beijing. All participants provided written informed consent, and the study was performed in accordance with the Declaration of Helsinki. The protocols for sample collection and use were approved by the Ethics Committee of the Beijing Institute of Genomics, Chinese Academy of Sciences.

### Droplet-based scRNA-seq and scATAC-seq using 10× Genomics chromium platform

We selected differentiated cells on days 7 and 14 from CB, PB, and SC during erythroid differentiation. The cell number and percent of living cells were counted using the Countstar Automated Cell Counter when the sample was used for cell capture, isolation, washing, and counting of nuclei suspensions, as per established protocols. Pre-indexed cells or nuclei were pooled and encapsulated in emulsion droplets using a standard microfluidic droplet generator (10× Genomics Chromium). The scRNA-seq and scATAC-seq technologies were used to obtain single-cell transcriptome data and chromatin accessibility. Briefly, the cells were loaded into each reaction for gel bead-in-emulsion (GEM) generation and cell barcoding. scRNA-seq and scATAC-seq libraries were constructed using the 10× Genomics Single-Cell 3’ Solution V3 kit and sequenced using an Illumina NovaSeq6000 sequencer.

### Integrated analysis of scRNA-seq and scATAC-seq

We calculated a gene activity score based on accessibility at a promoter region and the regulatory potential of nearby peaks and obtained a gene activity matrix from the scATAC-seq data. Specifically, gene coordinates for the human genome were obtained from EnsembleDB[49], and gene regions were extended to include a 2 kb upstream promoter region. Gene activities were assigned based on the number of fragments that mapped to each of the gene regions using the “FeatureMatrix” function, and gene activity scores were log normalized using the “NormalizeData” function in Signac[50] (v1.5.0). An anchoring approach was employed in the R Seurat59 (v4.1.1) and Signac packages to perform an integrative analysis of the scATAC-seq and scRNA-seq data. FindIntegrationAnchors and IntegrateData functions were called to combine the matrices. After integration, we ran RunPCA and performed UMAP embedding using runUMAP according to the standard Seurat workflow[51]. Clustering was performed using the FindNeighbors and FindClusters functions.

### Preliminary processing of scRNA-seq and scATAC-seq raw data from 10× genomics

Sequenced reads from scRNA-seq libraries were demultiplexed and aligned to the reference human genome GRCh38, barcode-processed, and unique molecular identifiers (UMI) were counted using the 10× Genomics Cell Ranger (v4.0.0) pipeline. Sequenced reads from the scATAC-seq libraries were processed using 10× Genomics Cell Ranger ATAC (v1.0.1). The sequencing reads were subjected to a Cell Ranger ATAC count pipeline consisting of read filtering, alignment to the reference genome using STAR[52], barcode counting, identification of transposase cut sites and accessible chromatin peaks, cell calls, and generation of count matrices for chromatin peaks and transcription factors.

### Cell type annotation

To annotate cell types based on scRNA-seq and scATAC-seq, we used a semi-supervised cell-type annotation method called SCINA[53] (v1.2.0). We combined cell type markers of hematopoietic cell types recorded in the PanglaoDB[54] and Atlas of Human Blood Cells database[55] to create a gene set for cell annotation. Unsupervised clustering was performed using the Seurat method, which does not rely on prior knowledge, and the same cluster generally contains the same cell type. At the same time, 10× genomics single-cell sequencing does not identify enough genes; hence, fluctuations in the expression of marker genes in a small number of cells are inevitable, resulting in false-positive cell annotation results. Therefore, we used the results of cell clusters as the main focus, which were complemented by the SCINA annotation results according to established hematopoietic cell markers, to count the percentage of cell types in each cell cluster and determine the cell type based on the principle of majority rule.

### Chromatin accessibility analysis

Analysis of chromatin datasets can be highly dependent on accurate peak calling and a bias in Cell Ranger for the aberrant merging of multiple distinct peaks into a single region[50]. To address this problem, we identified peaks using MACS2[56] (v2.0.3) for each annotated cell type separately and combined the individual peak calls into a unified peak set using Signac[50].

### Peak-to-gene linkage analysis

Advancements in single-cell sequencing technology have provided an opportunity to reconstruct gene regulatory networks (GRNs), including cis-regulatory elements relevant to regulation, by integrating scRNA-seq and scATAC-seq data. Non-coding elements can be linked to the regulation of gene expression through a correlation between DNA accessibility and the expression of nearby genes[57]. We used the peak-to-gene linkage method[58] in Signac. Briefly, we computed the Pearson correlation between the expression of a gene and accessibility of each peak within 500 kb of the gene transcriptional start site (TSS), and compared this value with the expected value given the GC content, overall accessibility, and length of the peak. Z-scores and p-values were calculated for each peak within the signal using the LinkPeaks function.

### Motif enrichment and TFs footprinting analysis

Motif analysis was performed using the chromVAR[59] R package (v1.16.0). In brief, we ran the AddMotifs function in Signac to add the DNA sequence motif information required for motif analysis. We then calculated a per-cell motif activity score by running chromVAR and identified the differential activity scores between the cell types.

From the JASPAR[60] database, we downloaded the degeneracy matrices of 633 TFs, which were input into FIMO[61] to search for hits. Highly accessible regions (HARs) were annotated with JASPAR motifs using the matchMotif option. The variability of JASPAR motifs across all cells was quantified using the computeDeviations function, followed by computing the variability using the computeVariability option. For each variable motif, we calculated mean z-scores for each cell type.

### Gene set variation analysis and functional enrichment analysis

Gene Ontology (GO) and Kyoto Encyclopedia of Genes and Genomes (KEGG) pathway analyses were performed using the clusterProfiler[62] (v4.2.2) package. The correlation analysis between Hif-1 and the cell cycle pathway was obtained using Pearson correlation analysis after calculating enrichment scores for pathways using the AddModuleScore function in the Seurat package. For gene set variation analysis, we used the R package GSVA (v1.42.0) with the Molecular Signatures Database[63] (MSigDB) hallmark gene sets to assess enrichment. GSVA transforms the gene expression data from an expression matrix of individual genes into an expression matrix of specific gene sets.

### Pseudo-time trajectory inference

To map the overall trajectory of cellular differentiation, we used the order_cells function in Monocle3 (v1.0.0) to compute pseudo-time values for each cell. We assessed hematopoietic stem and progenitor cells (HSPC) and erythroid cell developmental progression using Monocle2 (version 2.22.0). To order cells in pseudo-time, we applied the nonlinear dimensionality reduction algorithm DDRTree, which places cells along a trajectory corresponding to their biological differentiation states.

### Cell–cell communication analysis

CellChat[64] (v1.5.0), an R package described previously, contains ligand–receptor interaction databases for humans and mice that can analyze intercellular communication networks from scRNA-seq data annotated as different cell clusters. We used CellChat to assess global intercellular communication networks. We first constituted and quantified the global signaling cross-talk atlases, which provided the number of cell–cell interactions and their strengths in circle plots, and evaluated the major signaling inputs and outputs among all cell clusters using CellChatDB.human. We then used the netVisual_circle function to illustrate the strength or weakness of cell–cell communication networks from the target cell cluster to different cell clusters, and the netAnalysis_signalingRole_heatmap to identify signals contributing the most to *in vivo* or *in vitro* signaling. The netAnalysis_contribution function showed the contribution of significant ligand–receptor interactions between *in vivo* and *in vitro* signaling.

### Removing low-quality cells and doublets

For the scRNA-seq data analysis, low-quality cells were excluded based on a low number of genes detected (< 200) and/or high mitochondrial genetic content (> 20%). The DoubletFinder[65] (v2.0.3) package was used to identify doublets. For scATAC-seq data analysis, cells with sufficient numbers of fragments in peaks (> 3,500) and transcription start site (TSS) score ≥ 4 were kept. We calculated the doublet score using the ‘addDoubletScores’ function, and potential doublets were removed based on the ArchR[66] (v1.0.2).

### Flow cytometry analysis

Approximately 1–2 × 10^5^ cells were collected from the culture, centrifuged for 5 min at 300 × *g* to remove the supernatant, and washed twice with 1 ml 1× Dulbecco’s phosphate-buffered saline (DPBS; Catalog No. 14190136, Gibco) solution containing 2% fetal bovine serum (Catalog No. 16000044, Gibco) and 2 mM EDTA. After centrifugation, the cells were resuspended in 50 μL of 1× DPBS buffer, and the antibodies were added, followed by incubation at 4 °C for 10 min in the dark. The cells were washed twice by adding 1 mL of 1× DPBS buffer after incubation and resuspended in 200 μL of 1× DPBS buffer to prepare a cell suspension for the assay. A BD FACSAria II instrument was used for flow cytometry analysis, and data analysis was performed using the FlowJo software (v.7.6, Three Star). The antibodies used in this study included anti-human CD34-PE (Catalog No. 12-0349-41, eBioscience), anti-human CD45-FITC (Catalog No. 560976, BD Biosciences), anti-human CD71-PE (Catalog No. 130-099-219, Miltenyi Biotec), and anti-human CD235a-APC (Catalog No. 130-118-493, Miltenyi Biotec).

### Wright–Giemsa staining

Approximately 1 × 10^5^ cells were collected, and the medium was removed via centrifugation at 300 × *g* for 5 min. The cell pellets were resuspended in 1× DPBS. The cell concentration was adjusted to approximately 0.5 × 10^5^/mL. The slide was fixed on the rotating head of the slide ejector, and 0.2 mL cell suspension was added, centrifuged for 5 min at 1,000 rpm, and placed in a 37 °C incubator for 1–3 h. Then, 200 μL of Wright–Giemsa staining A solution (Catalog No. BA-4017, BASO, Zhuhai, China) was applied onto the cell-containing slide, and the slide was placed at room temperature for 1 min and covered with 600 μL of Wright–Giemsa staining B solution. After incubation at room temperature for 5 min, the slides were gently rinsed with distilled water along the area around the slides without cells for 1 min until the water became clear and transparent. The slide was placed at room temperature until dry, and the cell morphology was observed using a microscope.

## Acknowledgments

We are grateful to Prof. Linzhao Cheng for agreeing to use iPSC line BC1 and Prof. Qianfei Wang for providing the iPSCs. We are grateful for the expert assistance of Ms. Ting Li in the flow cytometry experiments and the core facility supporting data analysis from the China National Center for Bioinformation.

## Authorship Contributions

XF and ZZ conceived and supervised the study; ZX and GZ analyzed the data; WZ, LZ, ML, and TJ performed the experiments; LD facilitated the study design and contributed to the discussion; YS helped to isolate the HSPC from different sources; ZX, ZZ, and LZ wrote the manuscript; and XF, ZZ, and SH revised the manuscript.

## Data Availability

The raw sequence data reported in this article have been deposited in the Genome Sequence Archive[67] (GSA: Genome Sequence Archive) at the National Genomics Data Center, Beijing Institute of Genomics, Chinese Academy of Sciences/China National Center for Bioinformation (GSA: HRA005494), and are publicly accessible at https://ngdc.cncb.ac.cn/gsa-human/browse/HRA005494.

## Declarations

### Ethics approval and consent to participate

All participants provided written informed consent, and the study was performed in accordance with the Declaration of Helsinki. The protocols for sample collection and use were approved by the Ethics Committee of the Beijing Institute of Genomics, Chinese Academy of Sciences (No. 2018H020).

### Consent for publication

Not applicable

### Supplementary information

Additional file 1. Fig S1-S10

Additional file 2. Table S1. Marker genes of cell type (related to Figure 1)

Additional file 3. Table S2. Marker genes of erythroid cell subpopulations (related to Figure 4)

Additional file 4. Table S3. Enhancer-gene of different clusters and different systems (related to Figure 5)

Additional file 5. Table S4. Cell–cell communication of the erythroid differentiation system in vitro and in vivo (related to Figure 7)

---

**Figure S1.**
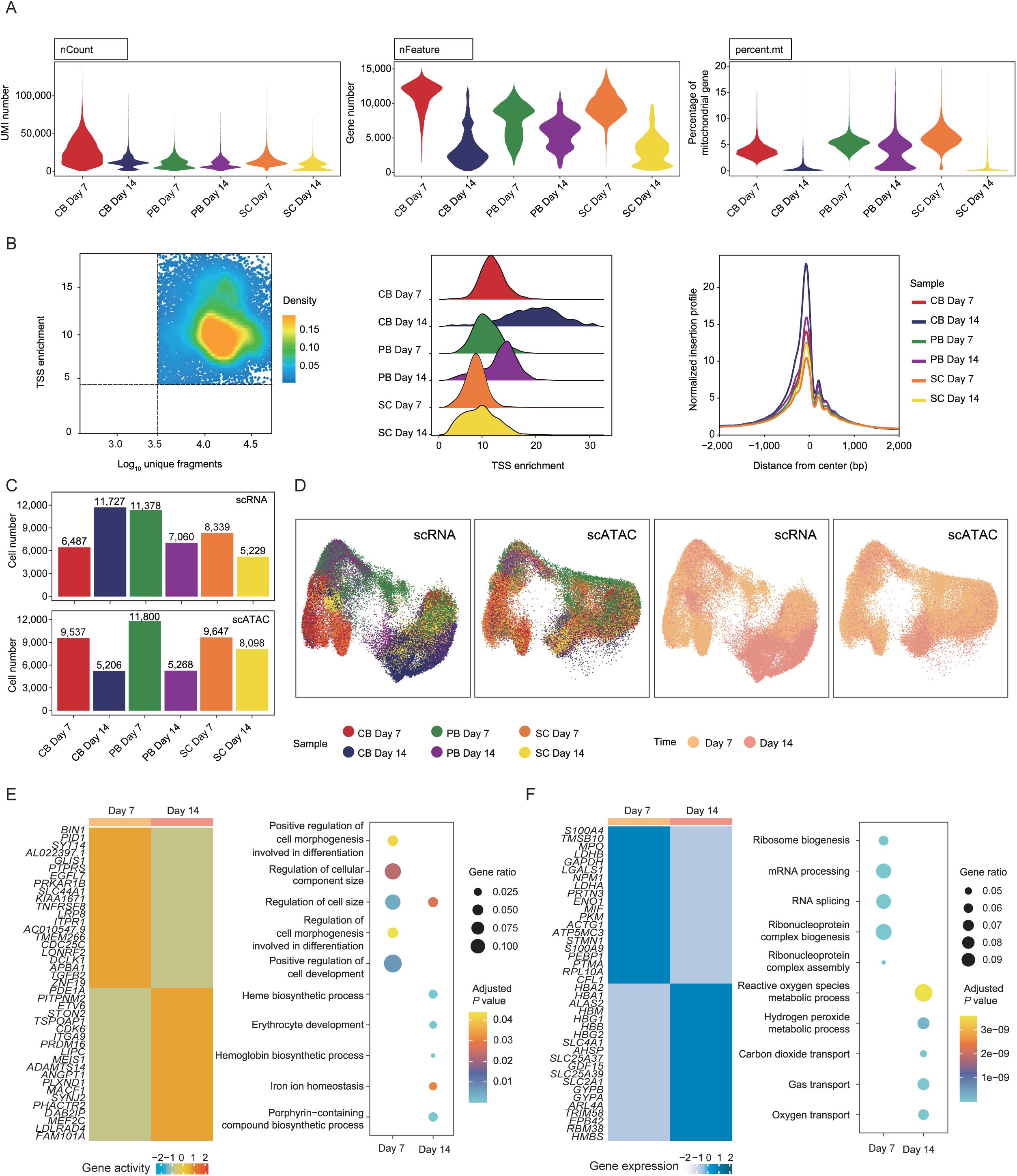

**Figure S2.**
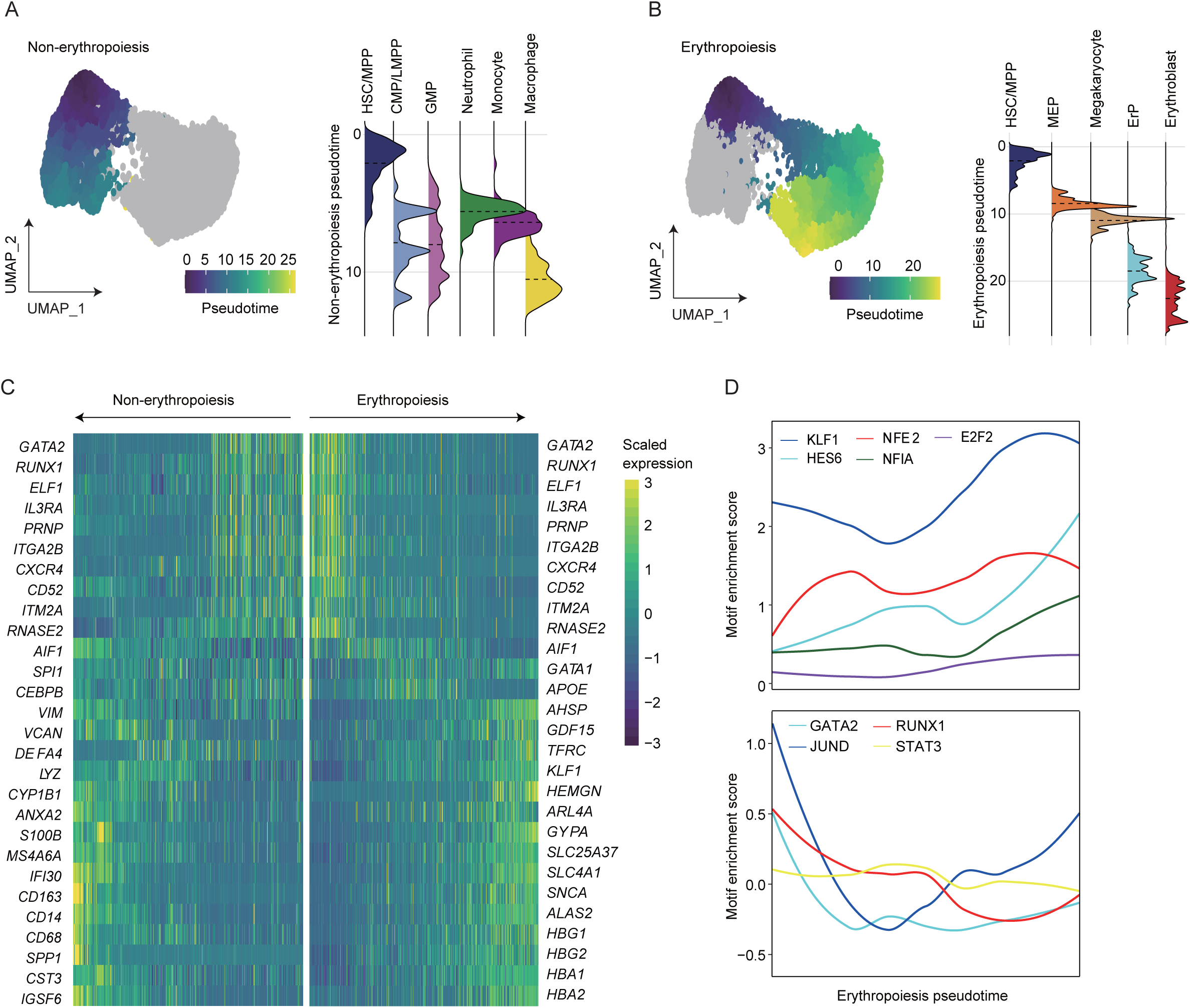

**Figure S3.**
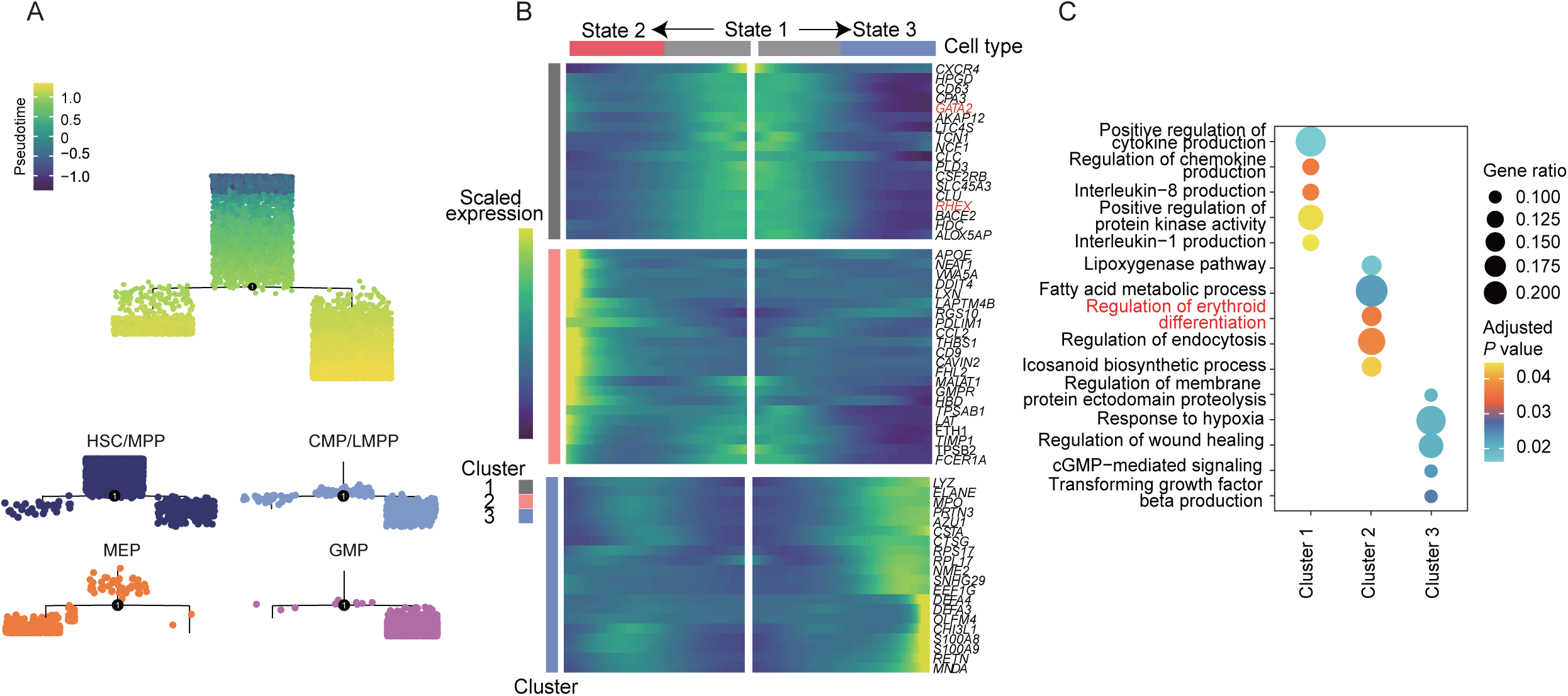

**Figure S4.**
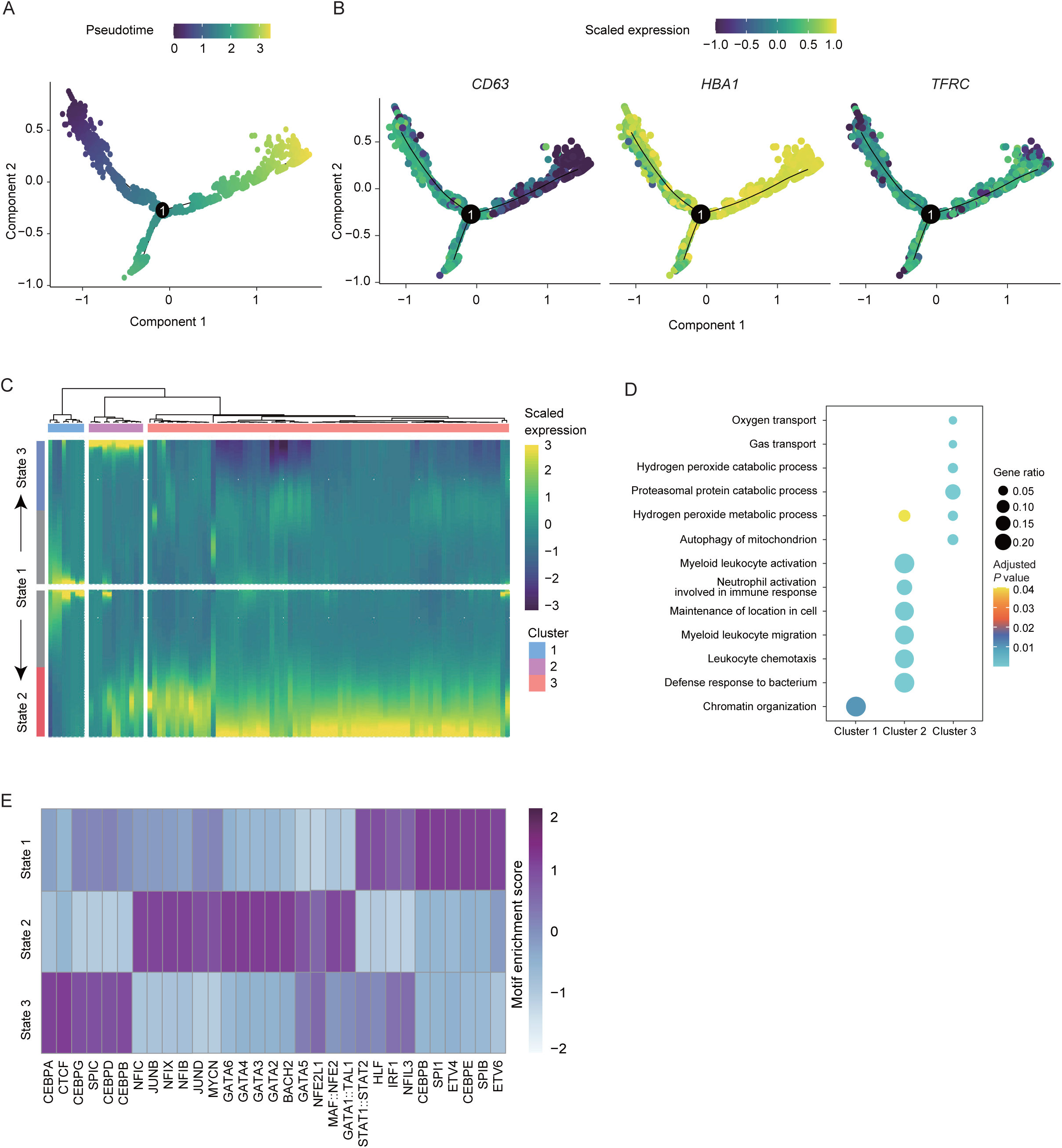

**Figure S5.**
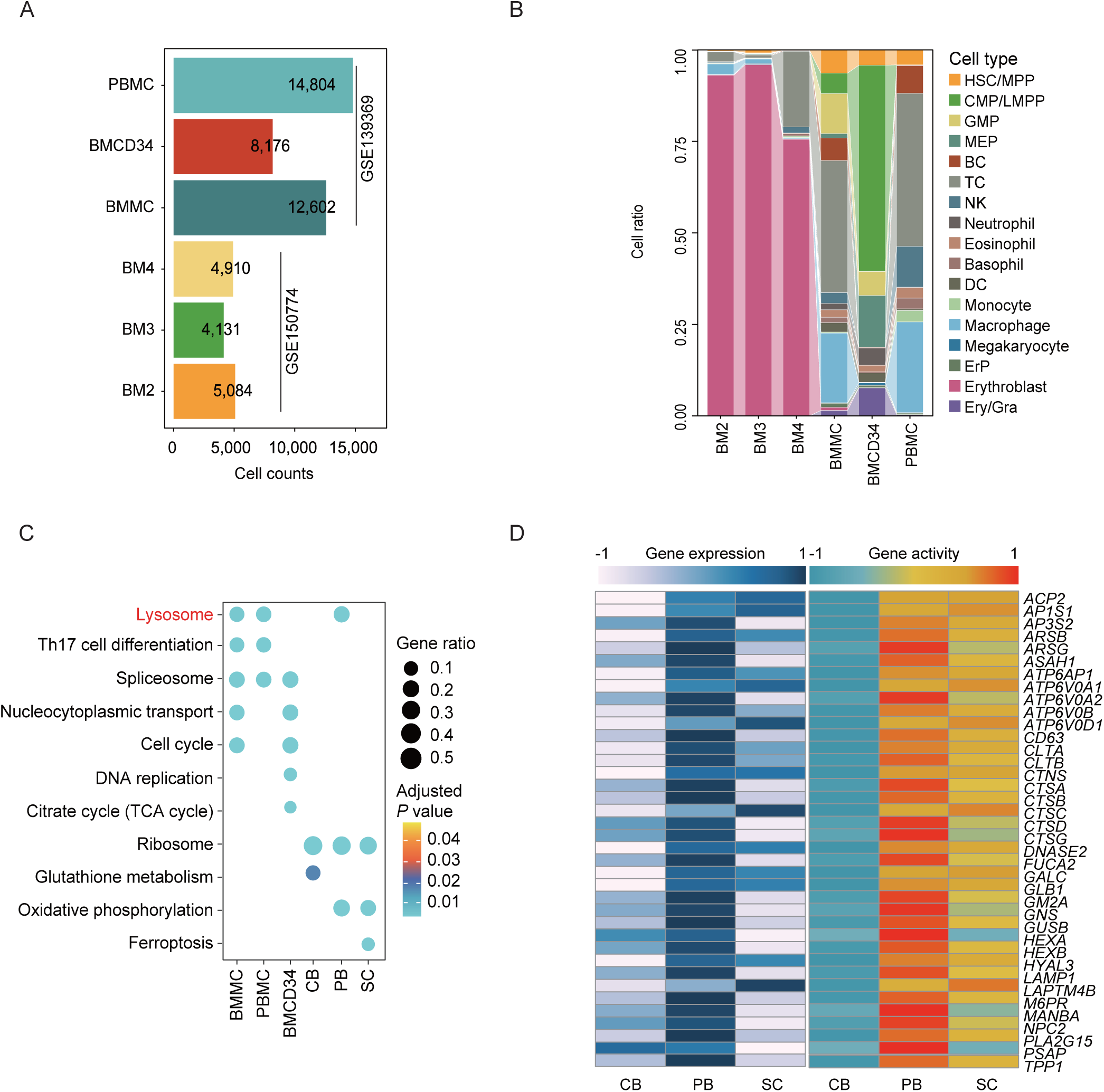

**Figure S6.**
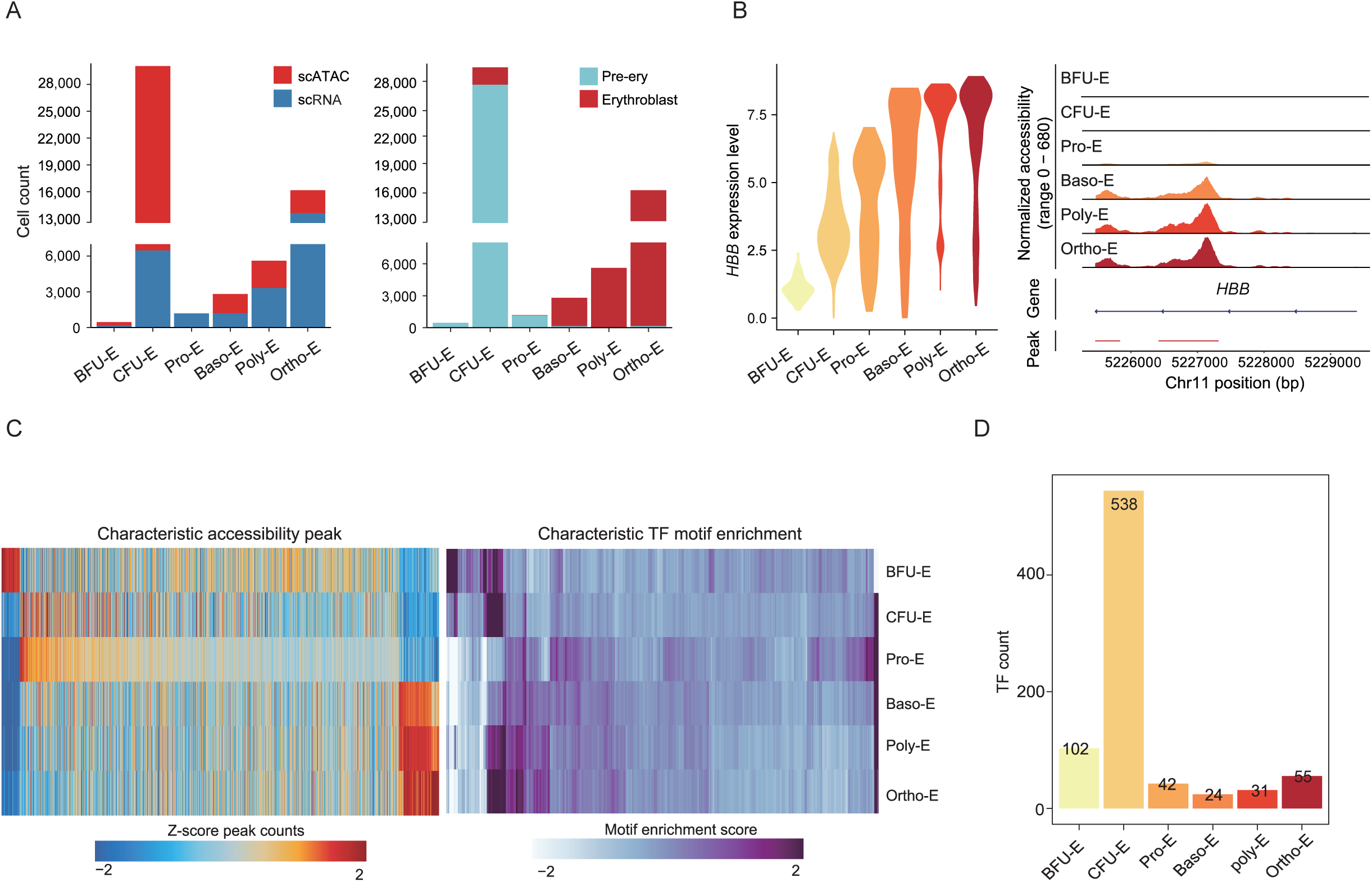

**Figure S7.**
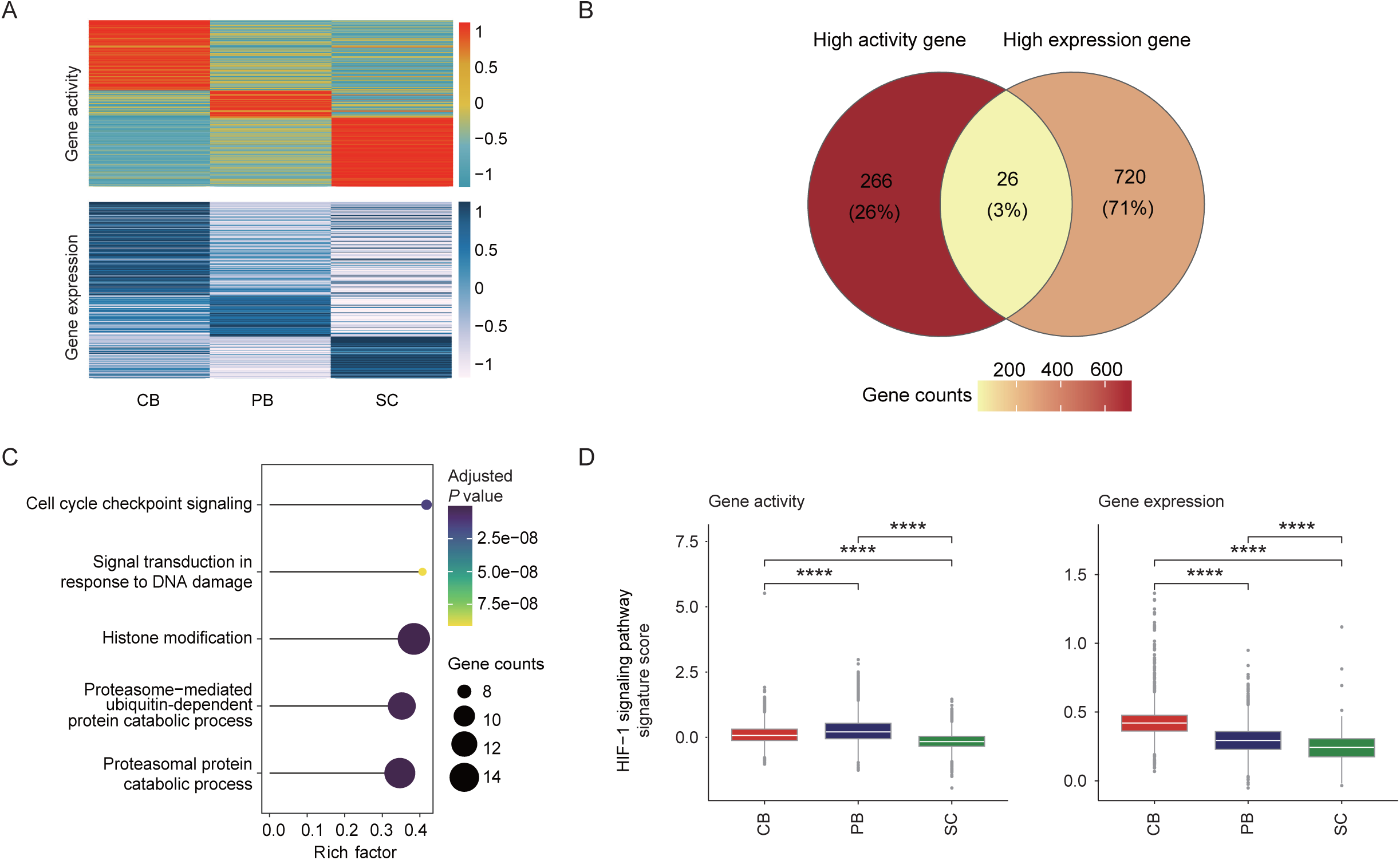

**Figure S8.**
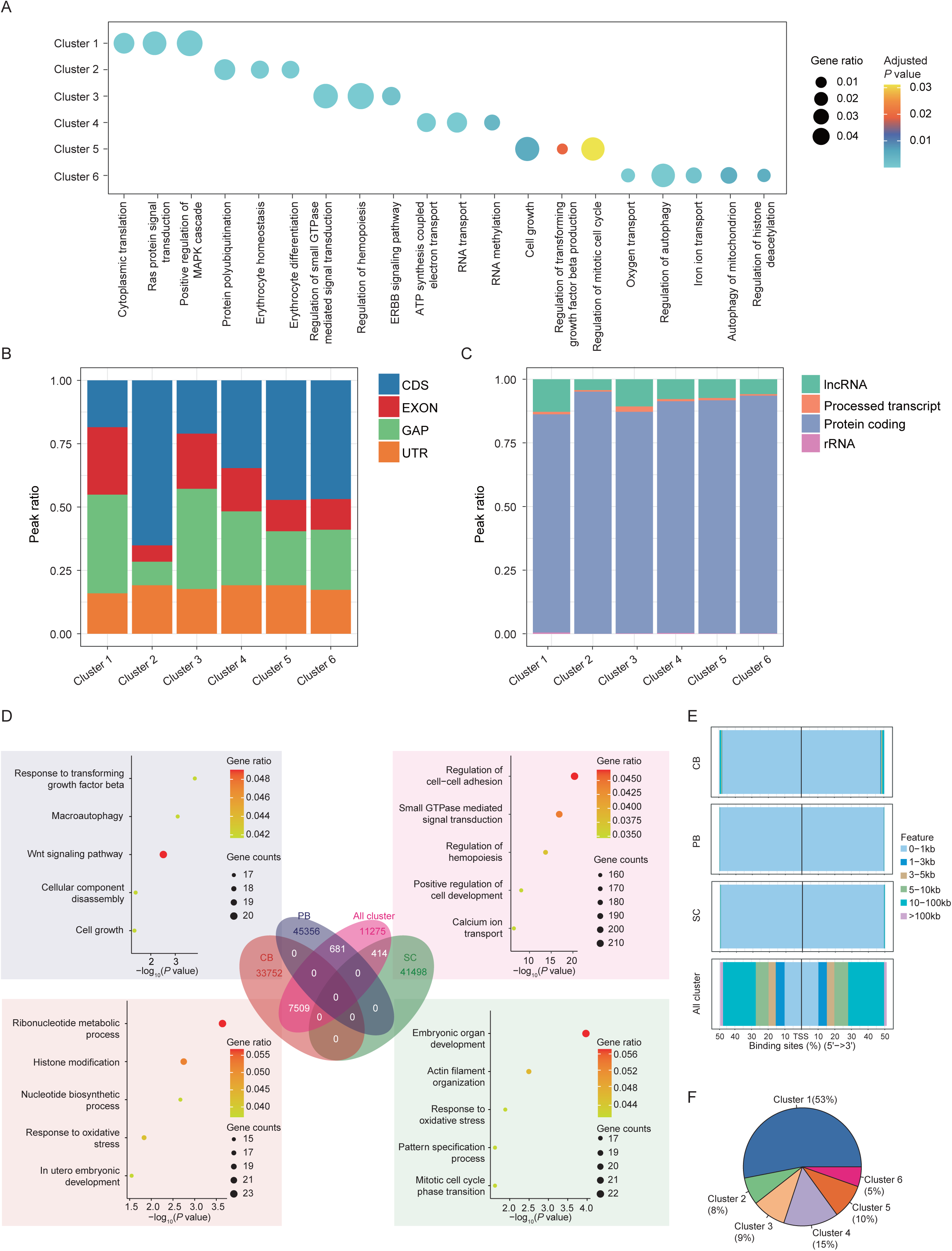

**Figure S9.**
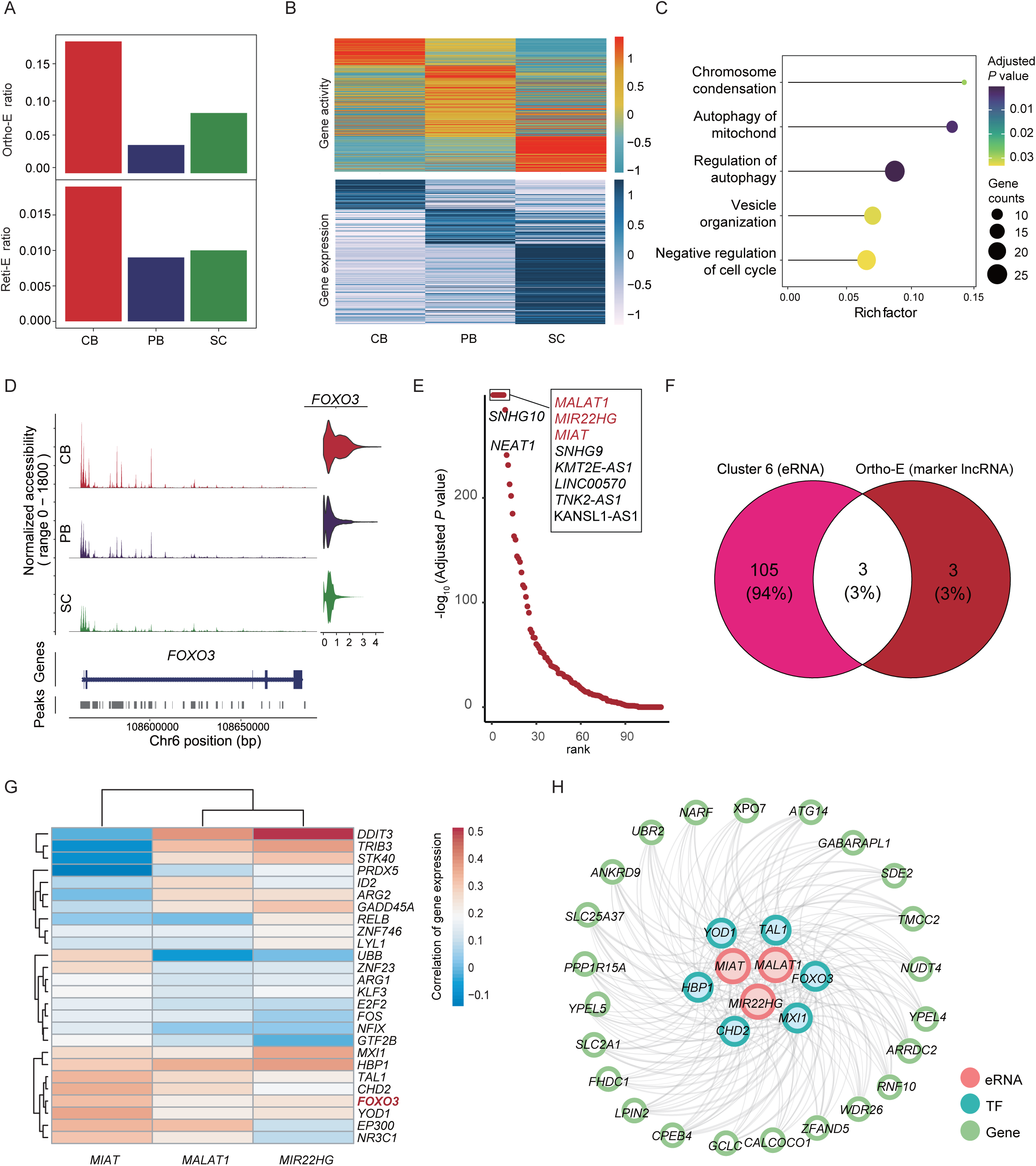

**Figure S10.**
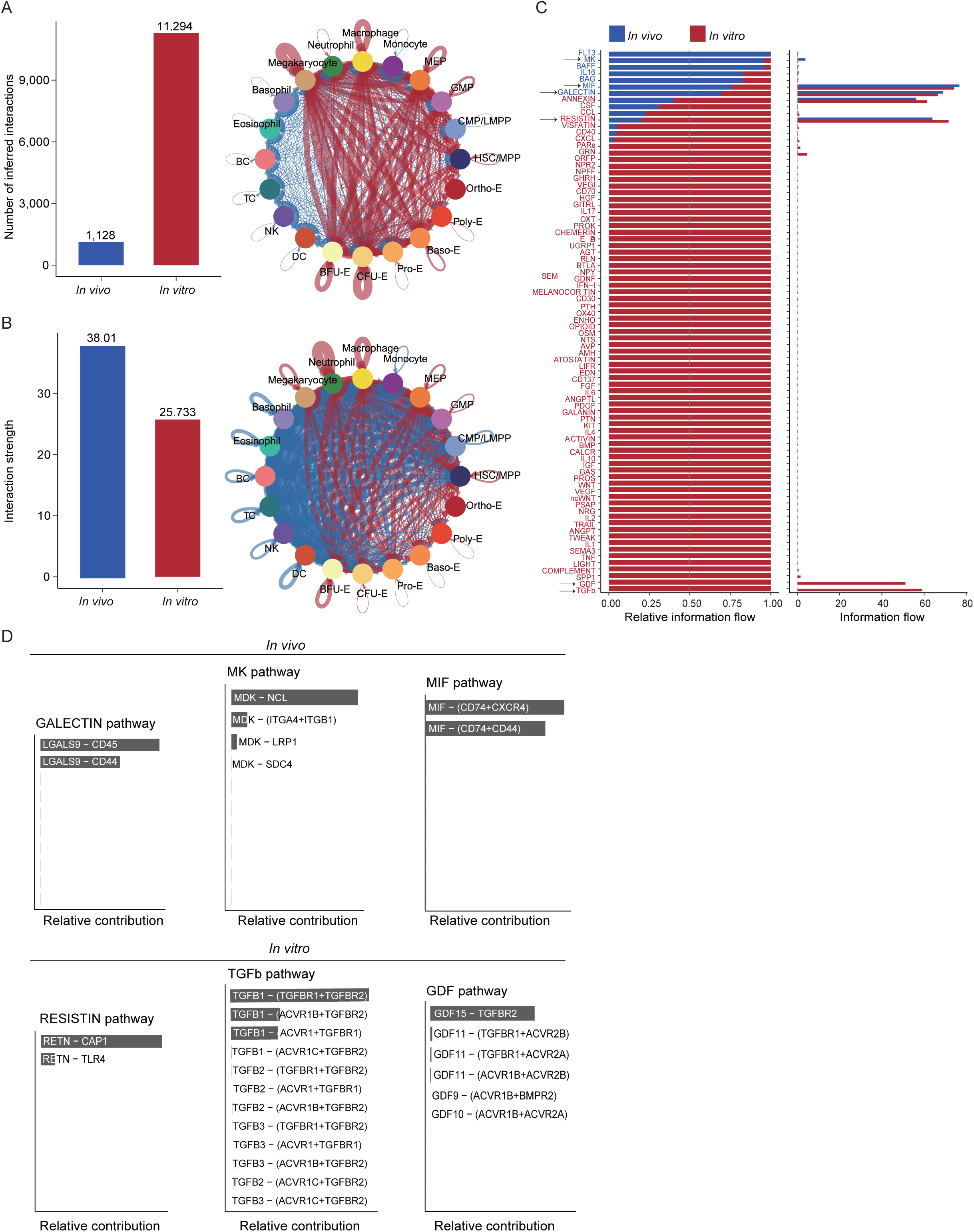

## References

1. Lee E, Sivalingam J, Lim ZR, Chia G, Shi LG, Roberts M, Loh YH, Reuveny S, Oh SKW: Review: generation of red blood cells for transfusion medicine: Progress, prospects and challenges. Biotechnol Adv 2018, 36:2118–2128.

2. Wang SH, Zhao HZ, Zhang H, Gao CJ, Guo XH, Chen LX, Lobo C, Yazdanbakhsh K, Zhang SJ, An XL: Analyses of erythropoiesis from embryonic stem cell-CD34(+) and cord blood-CD34(+) cells reveal mechanisms for defective expansion and enucleation of embryomic stem cell-erythroid cells. J Cell Mol Med 2022, 26:2404–2416.

3. Hansen M, von Lindern M, van den Akker E, Varga E: Human-induced pluripotent stem cell-derived blood products: state of the art and future directions. FEBS Letters 2019, 593:3288–3303.

4. Anstee DJ, Gampel A, Toye AM: Ex-vivo generation of human red cells for transfusion. Current Opinion in Hematology 2012, 19:163–169.

5. Wang XL, Zhang W, Zhao SQ, Yan H, Xin ZJ, Cui TT, Zang RG, Zhao LP, Wang HY, Zhou JN, et al: Decoding human in vitro terminal erythropoiesis originating from umbilical cord blood mononuclear cells and pluripotent stem cells. Cell Prolif 2024, 57:e13614.

6. Ludwig LS, Lareau CA, Bao EL, Nandakumar SK, Muus C, Ulirsch JC, Chowdhary K, Buenrostro JD, Mohandas N, An XL, et al: Transcriptional States and Chromatin Accessibility Underlying Human Erythropoiesis. Cell Reports 2019, 27:3228–3240 e3227.

7. Schippel N, Sharma S: Dynamics of human hematopoietic stem and progenitor cell differentiation to the erythroid lineage. Experimental Hematology 2023, 123:1–17.

8. Huang P, Zhao YZ, Zhong JM, Zhang XH, Liu QF, Qiu XX, Chen SK, Yan HX, Hillyer C, Mohandas N, et al: Putative regulators for the continuum of erythroid differentiation revealed by single -cell transcriptome of human BM and UCB cells. Proceedings of the National Academy of Sciences of the United States of America 2020, 117:12868–12876.

9. Jacobsen SEW, Nerlov C: Haematopoiesis in the era of advanced single-cell technologies. Nature Cell Biology 2019, 21:2–8.

10. Ye F, Huang WT, Guo GJ: Studying hematopoiesis using single-cell technologies. J Hematol Oncol 2017, 10:27.

11. Xin ZJ, Zhang W, Gong SJ, Zhu JW, Li YM, Zhang ZJ, Fang XD: Mapping Human Pluripotent Stem Cell-derived Erythroid Differentiation by Single-cell Transcriptome Analysis. Genomics Proteomics & Bioinformatics 2021, 19:358–376.

12. Oburoglu L, Tardito S, Fritz V, de Barros SC, Merida P, Craveiro M, Mamede J, Cretenet G, Mongellaz C, An X, et al: Glucose and glutamine metabolism regulate human hematopoietic stem cell lineage specification. Cell Stem Cell 2014, 15:169–184.

13. Li W, Wang Y, Zhao H, Zhang H, Xu Y, Wang S, Guo X, Huang Y, Zhang S, Han Y, et al: Identification and transcriptome analysis of erythroblastic island macrophages. Blood 2019, 134:480–491.

14. Leonard M, Brice M, Engel JD, Papayannopoulou T: Dynamics of Gata Transcription Factor Expression during Erythroid-Differentiation. Blood 1993, 82:1071–1079.

15. Verma R, Su S, McCrann DJ, Green JM, Leu K, Young PR, Schatz PJ, Silva JC, Stokes MP, Wojchowski DM: RHEX, a novel regulator of human erythroid progenitor cell expansion and erythroblast development. J Exp Med 2014, 211:1715–1722.

16. Xu CL, He J, Wang HT, Zhang YN, Wu J, Zhao L, Li Y, Gao J, Geng GF, Wang BR, et al: Single-cell transcriptomic analysis identifies an immune-prone population in erythroid precursors during human ontogenesis. Nat Immunol 2022, 23:1109–1120.

17. Arif T: Lysosomes and Their Role in Regulating the Metabolism of Hematopoietic Stem Cells. Biology-Basel 2022, 11:1410.

18. García-Prat L, Kaufmann KB, Schneiter F, Voisin V, Murison A, Chen J, Chan-Seng-Yue M, Gan OI, McLeod JL, Smith SA, et al: TFEB-mediated endolysosomal activity controls human hematopoietic stem cell fate. Cell Stem Cell 2021, 28:1838–1850.

19. Moon KR, van Dijk D, Wang Z, Gigante S, Burkhardt DB, Chen WS, Yim K, Elzen AVD, Hirn MJ, Coifman RR, et al: Visualizing structure and transitions in high-dimensional biological data. Nat Biotechnol 2019, 37:1482–1492.

20. Liu J, Zhang J, Ginzburg Y, Li H, Xue F, De Franceschi L, Chasis JA, Mohandas N, An X: Quantitative analysis of murine terminal erythroid differentiation in vivo: novel method to study normal and disordered erythropoiesis. Blood 2013, 121:e43–49.

21. Li J, Hale J, Bhagia P, Xue F, Chen L, Jaffray J, Yan H, Lane J, Gallagher PG, Mohandas N, et al: Isolation and transcriptome analyses of human erythroid progenitors: BFU-E and CFU-E. Blood 2014, 124:3636–3645.

22. Zhang FL, Shen GM, Liu XL, Wang F, Zhao YZ, Zhang JW: Hypoxia-inducible factor 1- mediated human GATA1 induction promotes erythroid differentiation under hypoxic conditions. Journal of Cellular and Molecular Medicine 2012, 16:1889–1899.

23. Smith BW, Rozelle SS, Leung A, Ubellacker J, Parks A, Nah SK, French D, Gadue P, Monti S, Chui DHK, et al: The aryl hydrocarbon receptor directs hematopoietic progenitor cell expansion and differentiation. Blood 2013, 122:376–385.

24. Pang CJ, Lemsaddek W, Alhashem YN, Bondzi C, Redmond LC, Ah-Son N, Dumur CI, Archer KJ, Haar JL, Lloyd JA, Trudel M: Kruppel-Like Factor 1 (KLF1), KLF2, and Myc Control a Regulatory Network Essential for Embryonic Erythropoiesis. Molecular and Cellular Biology 2012, 32:2628-2644.

25. Vinjamur DS, Wade KJ, Mohamad SF, Haar JL, Sawyer ST, Lloyd JA: Kruppel-like transcription factors KLF1 and KLF2 have unique and coordinate roles in regulating embryonic erythroid precursor maturation. Haematologica 2014, 99:1565–1573.

26. Tallack MR, Magor GW, Dartigues B, Sun L, Huang S, Fittock JM, Fry SV, Glazov EA, Bailey TL, Perkins AC: Novel roles for KLF1 in erythropoiesis revealed by mRNA-seq. Genome Res 2012, 22:2385–2398.

27. Liang R, Campreciós G, Kou Y, McGrath K, Nowak R, Catherman S, Bigarella CL, Rimmelé P, Zhang X, Gnanapragasam MN, et al: A Systems Approach Identifies Essential FOXO3 Functions at Key Steps of Terminal Erythropoiesis. PLoS Genet 2015, 11:e1005526.

28. Kumari R, Grzywa TM, Malecka-Gieldowska M, Tyszkowska K, Wrzesien R, Ciepiela O, Nowis D, Kazmierczak P: Ablation of Tmcc2 Gene Impairs Erythropoiesis in Mice. Int J Mol Sci 2022, 23:5263.

29. Lopez-Yrigoyen M, Yang CT, Fidanza A, Cassetta L, Taylor AH, McCahill A, Sellink E, von Lindern M, van den Akker E, Mountford JC, et al: Genetic programming of macrophages generates an in vitro model for the human erythroid island niche. Nature Communications 2019, 10:881.

30. Rhodes MM, Kopsombut P, Bondurant MC, Price JO, Koury MJ: Adherence to macrophages in erythroblastic islands enhances erythroblast proliferation and increases erythrocyte production by a different mechanism than erythropoietin. Blood 2008, 111:1700–1708.

31. Rabinovich GA, Vidal M: Galectins and microenvironmental niches during hematopoiesis. Current Opinion in Hematology 2011, 18:443–451.

32. Huang XL, Chao RH, Zhang YY, Wang PX, Gong XP, Liang DL, Wang YA: CAP1, a target of miR-144/451, negatively regulates erythroid differentiation and enucleation. J Cell Mol Med 2021, 25:2377–2389.

33. Olivier EN, Marenah L, McCahill A, Condie A, Cowan S, Mountford JC: High-Efficiency Serum-Free Feeder-Free Erythroid Differentiation of Human Pluripotent Stem Cells Using Small Molecules. Stem Cells Translational Medicine 2016, 5:1394–1405.

34. Romano L, Seu KG, Blanc L, Kalfa TA: Crosstalk between terminal erythropoiesis and granulopoiesis within their common niche: the erythromyeloblastic island. Current Opinion in Hematology 2023, 30:99–105.

35. Romano L, Seu KG, Papoin J, Muench DE, Konstantinidis D, Olsson A, Schlum K, Chetal K, Chasis JA, Mohandas N, et al: Erythroblastic islands foster granulopoiesis in parallel to terminal erythropoiesis. Blood 2022, 140:1621–1634.

36. Oburoglu L, Romano M, Taylor N, Kinet S: Metabolic regulation of hematopoietic stem cell commitment and erythroid differentiation. Current Opinion in Hematology 2016, 23:198–205.

37. Lyu J, Gu Z, Zhang Y, Vu HS, Lechauve C, Cai F, Cao H, Keith J, Brancaleoni V, Granata F, et al: A glutamine metabolic switch supports erythropoiesis. Science 2024, 386:eadh9215.

38. Noulin F, Manesia JK, Rosanas-Urgell A, Erhart A, Borlon C, Van Den Abbeele J, d’Alessandro U, Verfaillie CM: Hematopoietic Stem/Progenitor Cell Sources to Generate Reticulocytes for Culture. Plos One 2014, 9:e112496.

39. Rallapalli S, Guhathakurta S, Narayan S, Bishi DK, Balasubramanian V, Korrapati PS: Generation of clinical-grade red blood cells from human umbilical cord blood mononuclear cells. Cell Tissue Res 2019, 375:437–449.

40. Wang BR, Wang CC, Wan Y, Gao J, Ma YG, Zhang YN, Tong JY, Zhang YC, Liu JH, Chang LX, et al: Decoding the pathogenesis of Diamond-Blackfan anemia using single-cell RNA-seq. Cell Discovery 2022, 8:41.

41. Druker J, Wilson JW, Child F, Shakir D, Fasanya T, Rocha S: Role of Hypoxia in the Control of the Cell Cycle. Int J Mol Sci 2021, 22:4874.

42. Olivier E, Qiu CH, Bouhassira EE: Novel, High-Yield Red Blood Cell Production Methods from CD34-Positive Cells Derived from Human Embryonic Stem, Yolk Sac, Fetal Liver, Cord Blood, and Peripheral Blood. Stem Cells Translational Medicine 2012, 1:604–614.

43. Zhang JP, Yang ZX, Zhang F, Fu YW, Dai XY, Wen W, Zhang BL, Choi HN, Chen WQ, Brown M, et al: HDAC inhibitors improve CRISPR-mediated HDR editing efficiency in iPSCs. Science China-Life Sciences 2021, 64:1449–1462.

44. Smith DK, Yang JJ, Liu ML, Zhang CL: Small Molecules Modulate Chromatin Accessibility to Promote NEUROG2-Mediated Fibroblast-to-Neuron Reprogramming. Stem Cell Reports 2016, 7:955–969.

45. Jiang DF, Li TT, Guo CX, Tang TS, Liu HM: Small molecule modulators of chromatin remodeling: from neurodevelopment to neurodegeneration. Cell and Bioscience 2023, 13.

46. Tsiftsoglou AS, Vizirianakis IS, Strouboulis J: Erythropoiesis: model systems, molecular regulators, and developmental programs. IUBMB Life 2009, 61:800–830.

47. Chow A, Huggins M, Ahmed J, Hashimoto D, Lucas D, Kunisaki Y, Pinho S, Leboeuf M, Noizat C, van Rooijen N, et al: CD169+ macrophages provide a niche promoting erythropoiesis under homeostasis and stress. Nat Med 2013, 19:429–436.

48. Liu YF, Wang Y, Gao YX, Forbes JA, Qayyum R, Becker L, Cheng LZ, Wang ZZ: Efficient Generation of Megakaryocytes From Human Induced Pluripotent Stem Cells Using Food and Drug Administration-Approved Pharmacological Reagents. Stem Cells Translational Medicine 2015, 4:309–319.

49. Howe KL, Achuthan P, Allen J, Allen J, Alvarez-Jarreta J, Amode MR, Armean IM, Azov AG, Bennett R, Bhai J: Ensembl 2021. Nucleic acids research 2021, 49:D884–D891.

50. Stuart T, Srivastava A, Madad S, Lareau CA, Satija R: Single-cell chromatin state analysis with Signac. Nature methods 2021, 18:1333–1341.

51. Stuart T, Butler A, Hoffman P, Hafemeister C, Papalexi E, Mauck WM, 3rd, Hao Y, Stoeckius M, Smibert P, Satija R: Comprehensive Integration of Single-Cell Data. Cell 2019, 177:1888–1902 e1821.

52. Dobin A, Davis CA, Schlesinger F, Drenkow J, Zaleski C, Jha S, Batut P, Chaisson M, Gingeras TR: STAR: ultrafast universal RNA-seq aligner. Bioinformatics 2013, 29:15–21.

53. Zhang Z, Luo D, Zhong X, Choi JH, Ma Y, Wang S, Mahrt E, Guo W, Stawiski EW, Modrusan Z: SCINA: a semi-supervised subtyping algorithm of single cells and bulk samples. Genes 2019, 10:531.

54. Franzen O, Gan LM, Bjorkegren JLM: PanglaoDB: a web server for exploration of mouse and human single-cell RNA sequencing data. Database (Oxford) 2019, 2019:baz046.

55. Xie X, Liu M, Zhang Y, Wang B, Zhu C, Wang C, Li Q, Huo Y, Guo J, Xu C: Single-cell transcriptomic landscape of human blood cells. National Science Review 2021, 8:nwaa180.

56. Zhang Y, Liu T, Meyer CA, Eeckhoute J, Johnson DS, Bernstein BE, Nusbaum C, Myers RM, Brown M, Li W: Model-based analysis of ChIP-Seq (MACS). Genome biology 2008, 9:1–9.

57. Jansen IE, Savage JE, Watanabe K, Bryois J, Williams DM, Steinberg S, Sealock J, Karlsson IK, Hägg S, Athanasiu L: Genome-wide meta-analysis identifies new loci and functional pathways influencing Alzheimer’s disease risk. Nature genetics 2019, 51:404–413.

58. Ma S, Zhang B, LaFave LM, Earl AS, Chiang Z, Hu Y, Ding J, Brack A, Kartha VK, Tay T: Chromatin potential identified by shared single-cell profiling of RNA and chromatin. Cell 2020, 183:1103–1116. e1120.

59. Schep AN, Wu B, Buenrostro JD, Greenleaf WJ: chromVAR: inferring transcription-factor-associated accessibility from single-cell epigenomic data. Nature methods 2017, 14:975–978.

60. Fornes O, Castro-Mondragon JA, Khan A, Van der Lee R, Zhang X, Richmond PA, Modi BP, Correard S, Gheorghe M, Baranašić D: JASPAR 2020: update of the open-access database of transcription factor binding profiles. Nucleic acids research 2020, 48:D87–D92.

61. Bailey TL, Boden M, Buske FA, Frith M, Grant CE, Clementi L, Ren J, Li WW, Noble WS: MEME SUITE: tools for motif discovery and searching. Nucleic acids research 2009, 37:W202–W208.

62. Yu G, Wang LG, Han Y, He QY: clusterProfiler: an R package for comparing biological themes among gene clusters. OMICS 2012, 16:284–287.

63. Liberzon A, Subramanian A, Pinchback R, Thorvaldsdóttir H, Tamayo P, Mesirov JP: Molecular signatures database (MSigDB) 3.0. Bioinformatics 2011, 27:1739–1740.

64. Jin S, Guerrero-Juarez CF, Zhang L, Chang I, Ramos R, Kuan C-H, Myung P, Plikus MV, Nie Q: Inference and analysis of cell-cell communication using CellChat. Nature communications 2021, 12:1088.

65. McGinnis CS, Murrow LM, Gartner ZJ: DoubletFinder: doublet detection in single-cell RNA sequencing data using artificial nearest neighbors. Cell systems 2019, 8:329–337. e324.

66. Granja JM, Corces MR, Pierce SE, Bagdatli ST, Choudhry H, Chang HY, Greenleaf WJ: ArchR is a scalable software package for integrative single-cell chromatin accessibility analysis. Nature genetics 2021, 53:403–411.

67. Wang Y, Song F, Zhu J, Zhang S, Yang Y, Chen T, Tang B, Dong L, Ding N, Zhang Q: GSA: genome sequence archive. Genomics, proteomics & bioinformatics 2017, 15:14–18.

