## supplemental file1 for "Deciphering of single-cell chromatin accessibility and transcriptome reveals the discrepancy for *ex vivo* human erythropoiesis"

### Additional File 1

**Figure S1. Quality control of scRNA-seq and scATAC-seq data (related to Figure 1).**

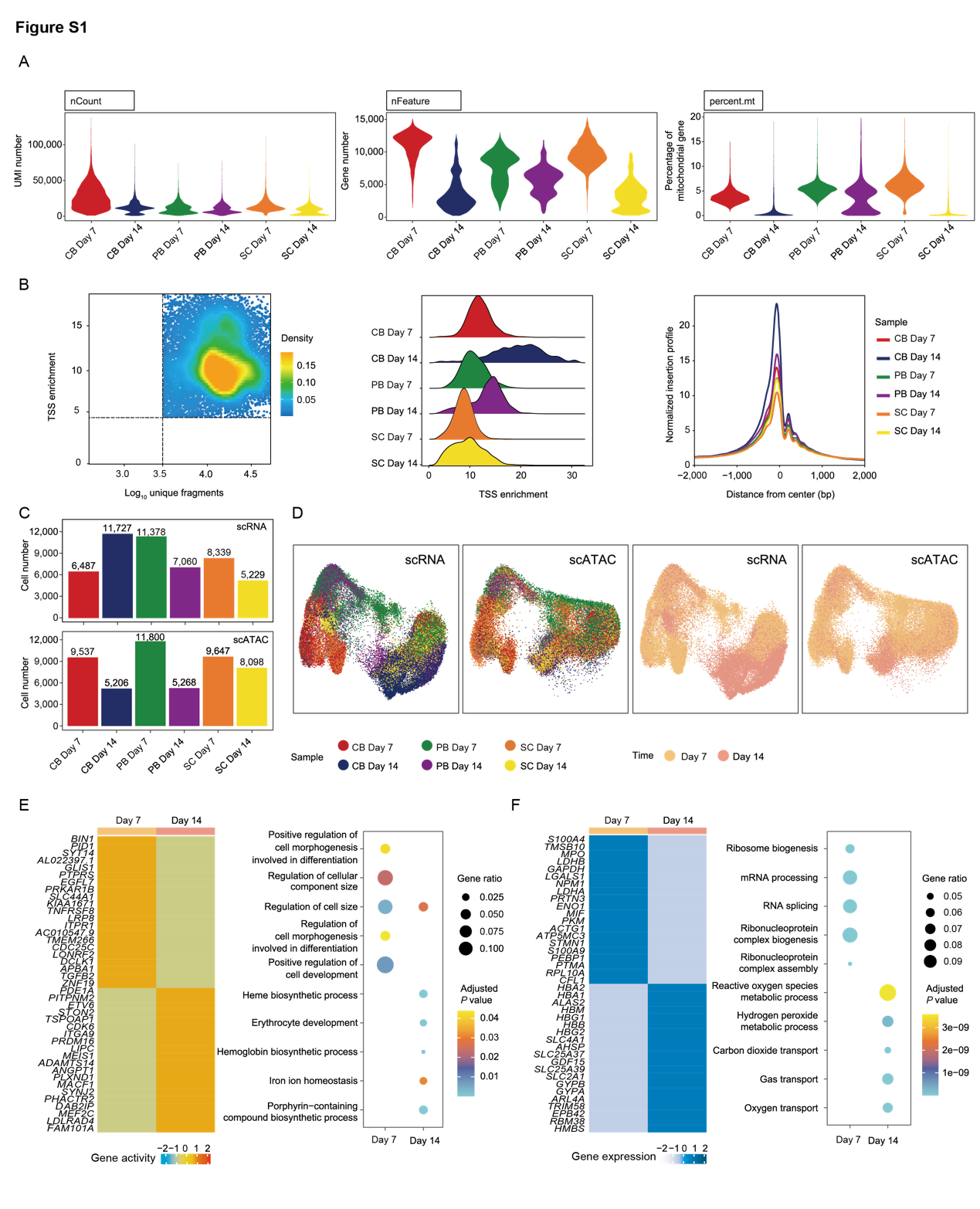

A. scRNA-seq data quality control results. Violin plots illustrating quality control results of each scRNA-seq sample, including the number of reads (nCount), number of genes (nFeature), and percentage of mitochondrial genes (percent.mt) per cell.
B. scATAC-seq data quality control results. Left, scatterplot showing the number of unique nuclear fragments and transcription start site (TSS) enrichment fraction per cell. Middle, ridge plot showing the TSS enrichment score of each sample. Right, line plot showing the TSS enrichment curve for each sample.
C. Bar plots showing the number of different cell types from scRNA-seq and scATAC-seq data separately.
D. The left two UMAP plots show the cell distribution at different systems with two differentiation states, and the right two UMAP plots show the cell distribution at two differentiation states based on scRNA-seq and scATAC-seq data.
E. Analysis of marker genes at different differentiation states of the scRNA-Seq data. The heatmap shows the top significant marker genes on Day 7 and Day 14 of the scRNA-seq data (left). The dot plot shows the functional annotation results of marker genes (right).
F. Analysis of marker genes at different differentiation states of the scATAC-Seq data. Heatmap showing the top significant annotated genes on Day 7 and Day 14 of the scATAC-seq data (left). Dot plot showing the functional annotation results of marker genes (right).

**Figure S2. Pseudo-time trajectory analysis of non-erythropoiesis and erythropoiesis in the *ex vivo* erythropoiesis system (related to Figure 1).**

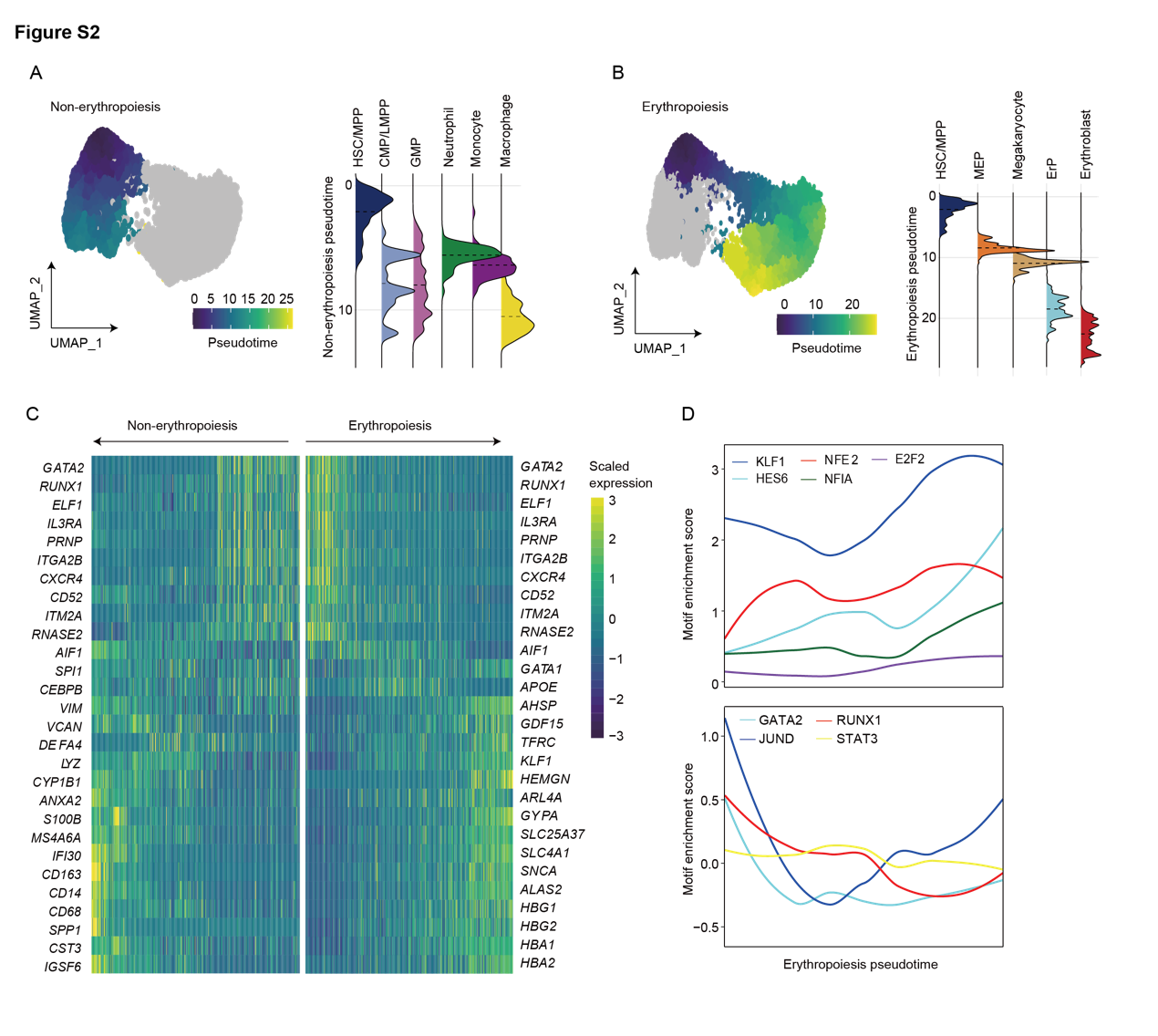

A. UMAP plots describing developmental pseudo-time trajectory from HSC/MPP to macrophages (Left, non-erythropoiesis). Abundance of non-erythropoiesis related cell types according to pseudo-time (Right).
B. UMAP plots describing developmental pseudo-time trajectory from HSC/MPP to erythropoiesis (Left, erythropoiesis). Abundance of erythropoiesis related cell types according to pseudo-time (Right).
C. Heatmap showing the differentially expressed genes along non-erythropoiesis and erythropoiesis branches.
D. Line plots showing enrichment scores of motifs that activate or repress erythropoiesis transcription according to pseudo-time trajectory.

**Figure S3. Early differentiation trajectory analysis of *ex vivo* erythropoiesis systems by using HSPC (related to Figure 2).**

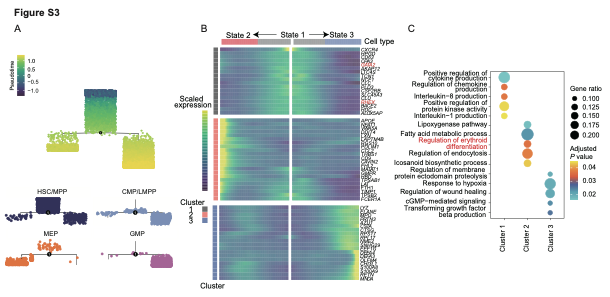

A. Pseudo-time differentiation trajectory of HSPC overlaid with each cell type separately (HSC/MPP, CMP/LMPP, MEP, and GMP).
B. Heatmap showing expression profiles of HSPC’s pseudo-time trajectory. The x-axis represents the timeline of trajectory analysis, the left y-axis represents the GO enrichment results’ cluster and the right y-axis represents the differentially expressed genes.
C. GO enriched terms of different clusters in Figure S3B’s left y-axis.

**Figure S4. Pseudo-time differentiation trajectory of erythroid cells in the *ex vivo* erythropoiesis system (related to Figure 2)**

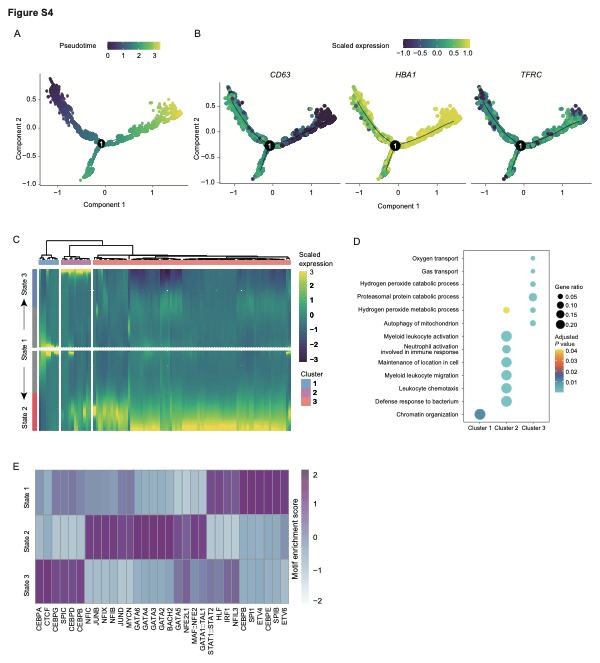

A. Pseudo-time differentiation trajectory of erythroid cells.
B. Expression patterns of *CD63*, *HBA1*, and *TFRC* in the pseudo-time differentiation trajectory of erythroid cells.
C. Heatmap showing expression profiles of erythroid cells pseudo-time trajectory.
D. GO enriched terms of different clusters in Figure S4C.
E. TF enrichment analysis of erythroid cells from different states in Figure S4C.

#### **Figure S5. Characterization of HSC in the *ex vivo* erythropoiesis system compared with that of *in vivo* sources**

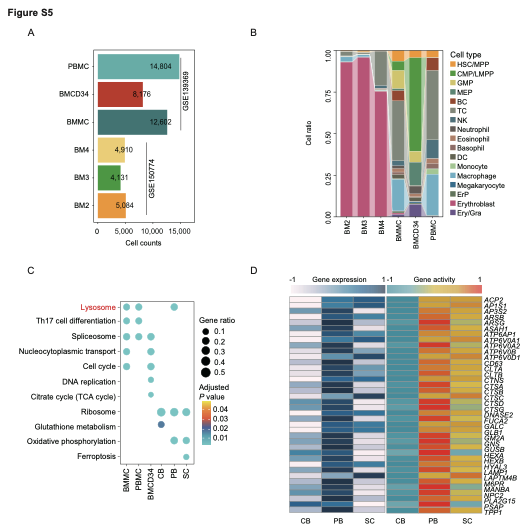

A. Bar plot describing the number of cell types from public scRNA-seq data *in vivo*.
B. Proportion of cell types from public scRNA-seq data *in vivo*.
C. GO enriched terms of HSC/MPP marker genes from different *ex vivo* erythropoiesis systems and public *in vivo* scRNA-seq data.
D. Heatmap showing the activity scores and expression levels of lysosome-related genes in state 1 cells derived from different erythropoiesis systems.

**Figure S6. Characteristics of each erythroid cell type in the ex vivo erythropoiesis systems (related to Figure 3).**

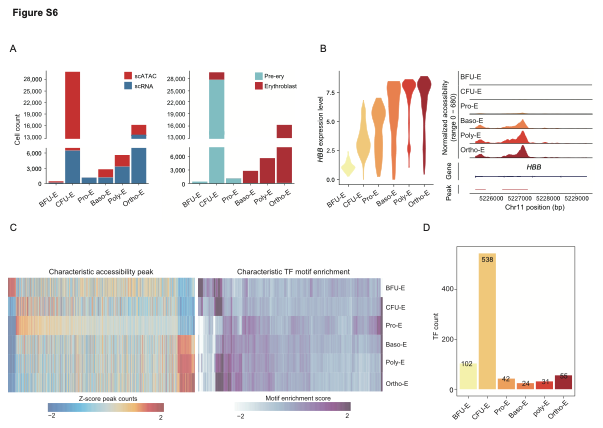

A. Bar plots illustrating the number of erythroid cell subpopulations colored by scRNA-seq and scATAC-seq data (left) or by ErP and erythroblast (right).
B. Gene expression and chromatin accessibility of *HBB* in each erythroid cell subpopulation. Left, Violin plot showing the gene expression level of *HBB* in each erythroid cell subpopulation; Right, chromatin accessibility peaks on *HBB* gene in each erythroid cell subpopulation. Normalized scATAC sequencing tracks of erythroid cell subpopulations at the *HBB* locus. Each track represents the average accessibility across each erythroid cell subpopulation.
C. Heatmaps showing the significant scATAC-seq accessibility peak count and enrichment score of TF’s motifs in each erythroid cell type.
D. Bar plot showing the TF motif count of erythroid cell type.

**Figure S7. Analysis of ErP differentiation defect in the *ex vivo* erythropoiesis systems (related to Figure 4).**

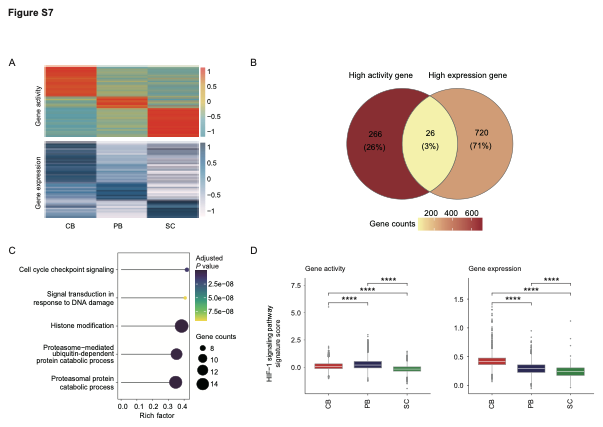

A. Heatmaps showing gene activity score and expression level of ErP (BFU-E and CFU-E) from different erythropoiesis systems.
B. Venn diagram showing the overlap between higher activity and more highly expressed genes of ErP in the CB-derived erythropoiesis system compared with those in the PB- and iPSC-derived erythropoiesis systems.
C. GO enriched terms of genes intersected in Figure S6B.
D. Comparison of HIF-1 signaling pathway signature’s scores of the gene activity score and expression level among different erythropoiesis systems. Statistical significance was determined by the two-side *Wilcoxon* rank-sum test. '****' *p* < 0.0001,'***' *p* < 0.001, '**' *p* < 0.01.

**Figure S8. Peak annotation in erythroid cells and enrichment results of marker peaks from three erythropoiesis systems (related to Figure 5).**

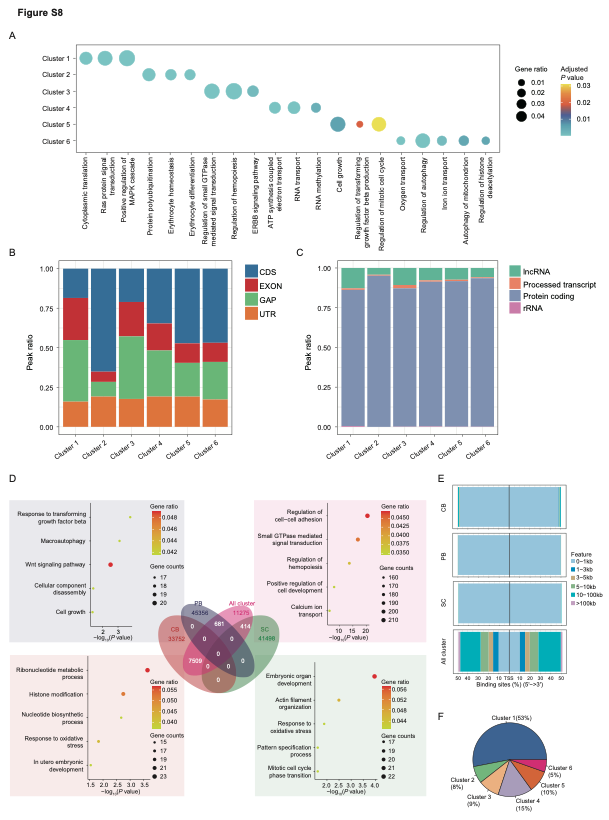

A. GO enriched terms of annotation genes from peaks in each cluster of Figure 5A.
B. The bar plot illustrating the distribution of genomic regions corresponding to peaks in each link cluster.
C. The bar plot showing the proportion of different RNA types corresponding to peaks in each link cluster
D. GO enrichment analysis and number of overlap peaks between all clusters in Figure 5B and marker peaks in each erythropoiesis system. Middle: Venn diagram showing the overlap between peaks of six clusters in erythroid cells and marker peaks in three erythropoiesis system. Bottom left: GO enrichment results for marker peaks in the CB-derived erythropoiesis system but not in six clusters. Bottom right: GO enrichment results for marker peaks in the SC-derived erythropoiesis system but not in six clusters. Top left: GO enrichment results for marker peaks in the PB-derived erythropoiesis system but not in six clusters. Top right: GO enrichment results at six clusters but not in any system marker peaks.
E. Proportion bars depicting the distribution of marker peaks’ binding sites in the CB-, PB-, and SC-derived systems, but not in six clusters and all clusters, and not in any system marker peaks.
F. Pie chart showing the percentage of each cluster’s peak but not in any *ex vivo* erythropoiesis system.

**Figure S9. Erythroid enucleation discripancy across three ex vivo erythropoiesis systems.**

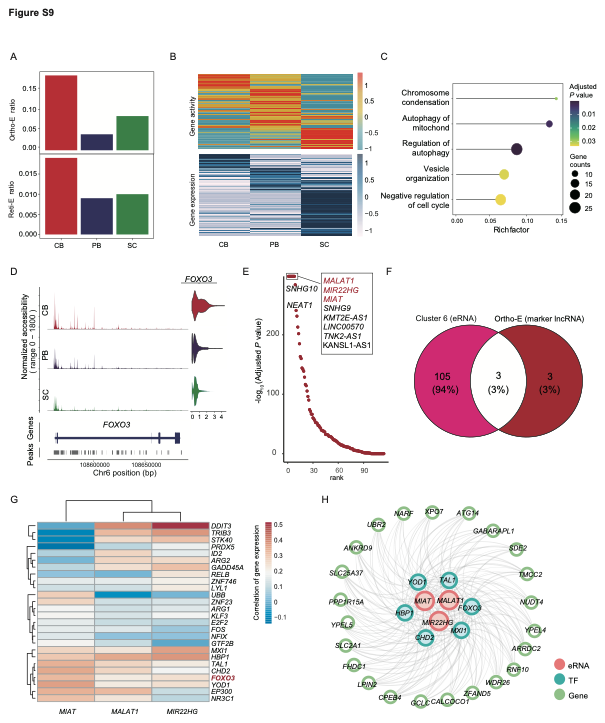

A. Cell number ratios of Ortho-E and Reti-E from different erythropoiesis systems.
B. Heatmap showing the gene activity score and expression level of Ortho-E in different erythropoiesis systems.
C. GO-enriched terms of highly expressed genes in Ortho-E derived from the CB-derived system.
D. Chromatin accessibility peaks and gene expression level of *FOXO3* in Ortho-E cells from different *ex vivo* erythropoiesis systems.
E. Dot plot showing significant lncRNAs identified in Ortho-E of the CB-derived erythropoiesis system.
F. Venn diagram showing the overlap between the significant IncRNAs in Ortho-E of the CB-derived system and the potential enhancer peaks of cluster 6 in Figure 5B.
G. Heatmap showing the correlation results between the common eRNA in Figure S9H and the highly expressed TF-encoding gene of Ortho-E in the CB-derived erythropoiesis system.
H. Regulatory network of eRNAs, TFs, and genes in Ortho-E in the CB-derived erythropoiesis system.

**Figure S10. Inferred communication networks between *in vivo* and *in vitro* differentiation systems (related to Figure 7).**

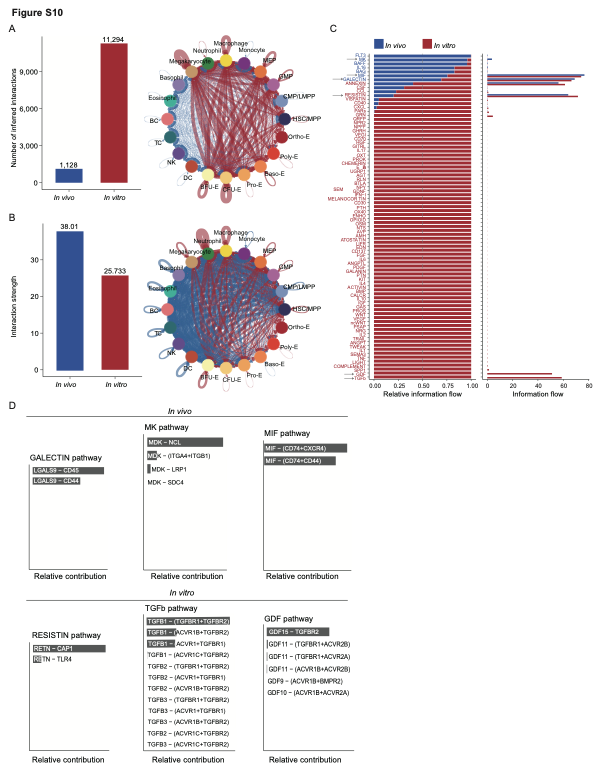

A. Comparison of the number of inferred interactions between *in vivo* and *in vitro*.
B. Comparison of the strength of inferred interactions between *in vivo* and *in vitro*.
C. Ranking signaling networks based on the information flow. Significant signaling pathways were ranked based on differences in the overall information flow within the inferred networks between *in vivo* and *in vitro*.
D. Relative contribution of ligand–receptor pairs in the GALECTIN, MK, and MIF signaling pathways *in vivo*, and in the RESISTIN, TGFβ, and DGF signaling pathways *in vitro*.
