## supplemental file2 for "Deciphering of single-cell chromatin accessibility and transcriptome reveals the discrepancy for *ex vivo* human erythropoiesis"

**Additional File 2**

**Table S1. Marker genes of cell type (related to Figure 1)**

The SHEET "*Markers from scRNA*" shows markers for cell populations from scRNA-seq data.

The SHEET "*Markers from scATAC*" shows markers for cell populations from scATAC-seq data.

**Table S2. Marker genes of erythroid cell subpopulations (related to Figure 3 and Figure S9)**

The SHEET *"Markers of IncRNA"* shows marker genes for erythroid cell subpopulations of lncRNA.

The SHEET *"Markers of protein"* shows marker genes for erythroid cell subpopulations of protein.

The SHEET *"Markers of TF"* shows marker genes for erythroid cell subpopulations of TF.

**Table S3. Enhancer-gene of different clusters and different systems (related to Figure 5 and Figure S9)**

The SHEET *"Peaks of clusters"* shows peaks of erythroid cell subpopulations from cluster 1 to 6.

The SHEET *"Peaks of systems"* shows peaks of erythroid cell subpopulations from three differentiation system (SC, PB, and CB).

**Table S4. Cell–cell communication of the erythroid differentiation system in vitro and in vivo (related to Figure 7)**

The SHEET *"Pathways in vitro"* shows cell–cell communication pathways of the erythroid differentiation system *in vitro.*

The SHEET *"Pathways in vivo"* shows cell–cell communication pathways of the erythroid differentiation system *in vivo.*
